# Global transcriptomics reveals specialized roles for splicing regulatory proteins in the macrophage innate immune response

**DOI:** 10.1101/2020.12.06.413690

**Authors:** KO West, AR Wagner, HM Scott, KJ Vail, KE Carter, RO Watson, KL Patrick

## Abstract

Pathogen sensing via pattern recognition receptors triggers massive reprogramming of macrophage gene expression. While the signaling cascades and transcription factors that activate these responses are well-known, the role of post-transcriptional RNA processing in modulating innate immune gene expression remains understudied. Recent phosphoproteomics analyses revealed that members of the SR and hnRNP families of splicing regulatory proteins are dynamically post-translationally modified in infected macrophages. To begin to test if these splicing factors play a privileged role in controlling the innate immune transcriptome, we analyzed steady state gene expression and alternatively spliced isoform production in ten SR/hnRNP knockdown RAW 264.7 macrophage cell lines following infection with the bacterial pathogen *Salmonella enterica* serovar Typhimurium (*Salmonella*). We identified thousands of transcripts whose abundance was increased or decreased by SR/hnRNP knockdown in macrophages. We observed that different SR/hnRNPs control the expression of distinct gene regulons in uninfected and *Salmonella*-infected macrophages, with several key innate immune genes (*Nos2, Mx1, Il1a*) relying on multiple SR/hnRNPs to maintain proper induction and/or repression. Knockdown of SR/hnRNPs promoted differential isoform usage (DIU) for a number of key immune sensors and signaling molecules and many of these splicing changes were again, distinct in uninfected and *Salmonella*-infected macrophages. Finally, after observing a surprising degree of similarity between the DEGs and DIUs in hnRNP K and U knockdown macrophages, we found that these cells are better able to restrict vesicular stomatitis virus replication than control cells, supporting a role for these hnRNPs in controlling infection outcomes in macrophages *ex vivo*. Based on these findings, we conclude that many innate immune genes have evolved to rely on one or more splicing regulatory factors to ensure the proper timing and magnitude of their induction, bolstering a model wherein pre-mRNA splicing is a critical regulatory node in the innate immune response.

## INTRODUCTION

When innate immune cells like macrophages sense pathogens, they undergo dramatic gene expression reprogramming and upregulate thousands of genes. Proper regulation of the timing and magnitude of innate immune gene induction is critical to ensure that the immune sys-tem is adequately engaged to fend off microbial invaders without risking deleterious outcomes associated with hyperinflammation [1-3]. While there has been great interest in the nature of macrophage sensing and signaling events that activate a transcriptional response following an inflammatory signal, much less is known about how regulation of RNA processing steps down-stream of transcription influence innate immune gene expression outcomes.

Consistent with the current “transcription-focused” paradigm of innate immune gene expression, research has categorized innate immune genes into primary and secondary response genes [4, 5]. Primary, or early response genes, are readily induced upon activation of pathogen sensing cascades. Many of these transcripts reach maximal abundance as soon as 30 minutes post-pathogen sensing [6, 7], whereas secondary response genes (the activation of which requires nascent protein synthesis of a transcription factor or expression of a cytokine to activate another wave of gene expression via autocrine signaling), are maximally induced with slower kinetics. The timing and induction of primary and secondary response genes relies on a number of tightly regulated mechanisms, including but not limited to, cooperative binding of transcription factors [8, 9], nucleosome occupancy and histone modification at promoters [10, 11], signal dependent interactions between transcription factor subunits [12, 13], and selective interaction with transcriptional elongation machinery [14].

On top of this intricate transcriptional regulatory network, once innate immune transcripts are transcribed, they, like most eukaryotic transcripts, are subject to post-transcriptional regulation at the level of pre-mRNA splicing, cleavage and polyadenylation, mRNA export, and non-sense mediated decay. Pre-mRNA splicing is being increasingly appreciated as an important regulatory node in cells undergoing stress or responding to extracellular triggers, including exposure to vitamins and metal ions [15], heat shock [16, 17], and UV damage [18, 19]. There is growing interest in how RNA processing modulates innate immune gene expression outcomes in macrophages. Both *Salmonella enterica* and *Listeria monocytogenes* infection promote widespread 3’UTR shortening and exon inclusion in primary human macrophages [20] and alternative splicing and nonsense mediated decay play an important role in balancing isoform abundance of key antiviral innate immune molecules like *Oas1g* [21]. Important kinetic studies of gene expression in Lipid A-treated bone marrow derived macrophages showed that removal of introns from pre-mRNAs and release of processed innate immune transcripts from chromatin can be significantly delayed relative to onset of a gene’s transcription [6, 7]. These data are consistent with numerous reports over the last decade demonstrating that the majority of splicing, both intron recognition and catalysis, occurs co-transcriptionally [22-25]. While these findings argue that post-transcription regulatory mechanisms play a key role in controlling the timing and abundance of translation-competent mRNAs and thus directly influence the overall production of cytokines, chemokines, and antimicrobial proteins, we still know very little about the mechanisms that drive this regulation and the specific macrophage proteins involved.

One mechanism through which splicing factors may be regulated during the innate immune response is by receiving signals from outside the cell, for example, via phosphorylation of SR family proteins [16, 17, 26]. Defined on basis of possessing a serine-arginine rich motif, SR proteins play a key role in directing the spliceosome to particular regions of a transcript by binding exonic splicing enhancers. SR proteins often work cooperatively and antagonistically with proteins in the heterogenous nuclear ribonucleoprotein (hnRNP) family, which also bind to conserved sequences in exons and introns to influence splicing decisions. We became interested in a role for SR and hnRNP proteins in regulating macrophage gene expression after several global phospho-proteomics studies revealed that proteins involved in mRNA processing are among the most differentially phosphorylated proteins in macrophages following infection with a bacterial [27, 28] or fungal pathogen [29], despite showing no significant change in protein abundance. These data led us to speculate that splicing regulatory proteins are functionalized following pathogen sensing.

To begin to define the contribution of splicing regulatory proteins to modulating innate immune gene expression, we took an unbiased approach and knocked down expression of ten members of the SR/hnRNP families of splicing regulatory factors. We infected these knockdown cell lines with *Salmonella enterica* serovar Typhimurium (*Salmonella*) and measured differential gene expression (DGE) and differential isoform usage (DIU) in steady state RNA at a key innate immune time point. Our analysis found that most SR/hnRNPs regulate the abundance or splicing of different cohorts of genes and most that SR/hnRNP-sensitive DEGs are not also subject to DIU. While the reliance of innate immune transcripts on SR/hnRNPs did not correlate with induction level, gene length, or exon/intron number, we did observe that many rapidly induced “primary response” genes are hyper-induced in the SR/hnRNP knockdown macrophages following *Salmonella* infection, suggesting that pre-mRNA splicing helps repress or limit induction of these transcripts. Together, our data implicate splicing regulatory protein networks in fine-tuning the timing and magnitude of the induction of a cohort of innate immune transcripts and highlight a previously unappreciated role for RNA binding proteins and post-transcriptional control mechanisms in regulating macrophage gene expression and antimicrobial capacity.

## RESULTS

### SR and hnRNPs control the abundance of distinct sets of transcripts in uninfected and *Salmonella*-infected macrophages

To understand how splicing regulatory proteins shape global innate immune gene expression, we sought to identify factors most likely to play a privileged role in the macrophage innate immune response. Two recent publications identified a number of splicing factors that were differentially phosphorylated during bacterial infection (specifically during infection with the intracellular bacterial pathogen *Mycobacterium tuberculosis*) in a RAW 264.7 macrophage cell line [27] or in primary mouse macrophages [28]. We manually annotated these lists and prioritized a set of SRs and hnRNPs for transcriptomics analysis. Based on these proteomics data, SRSF1, SRSF2, SRSF6, SRSF7, SRSF9, hnRNP C, hnRNP F, hnRNP K, hnRNP M, and hnRNP U were predicted to be phosphorylated or dephosphorylated at one or more amino acid residues over the first 24h of *M. tuberculosis* infection (Fig. S1A). To begin to determine how each of these factors contribute to gene expression in macrophages, we generated murine macrophage cell lines (RAW 264.7) in which each factor was stably knocked down via expression of an shRNA construct targeting either an exon or the 3’UTR for each factor, with regions chosen to ensure that all protein coding isoforms of each factor would be targeted by the shRNA (Fig. 1A). A notable caveat of these stable knockdown cell lines is that overall knockdown efficiency varied between factors, with only about 50% knockdown efficiency achieved for hnRNP C, hnRNP K, SRSF1 and SRSF7 and 70-90% knockdown achieved for SRSF2, SRSF6, SRSF9, hnRNP F and hnRNP U. We predict this variation in knockdown efficiency is indicative of the macrophage’s ability to tolerate loss of each of these factors and likely correlates with the cell’s reliance on each for proper RNA processing of essential housekeeping genes.

**Figure 1:**
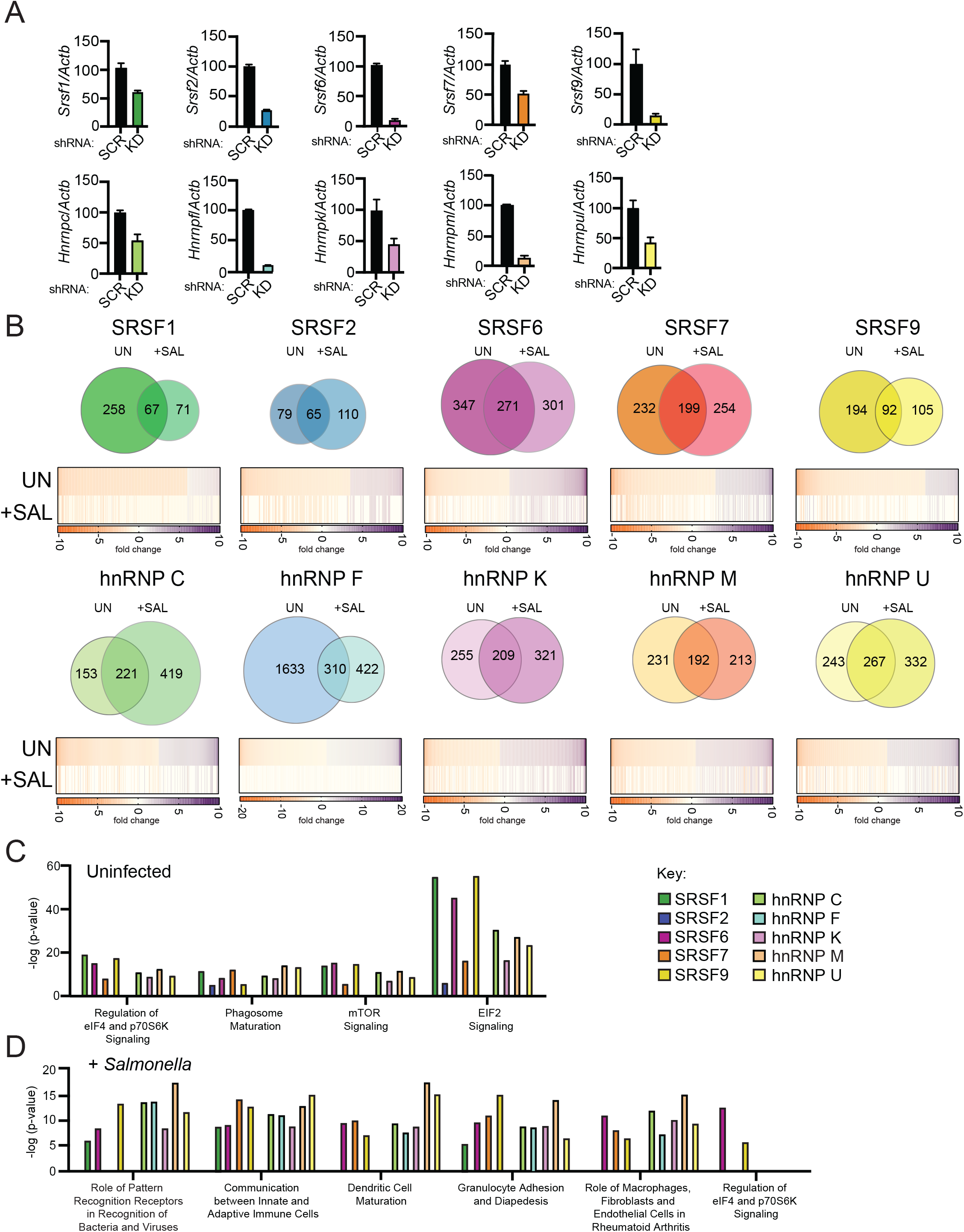
RNA-Seq reveals distinct hnRNP- and SRSF-dependent regulons in uninfected and *Salmonella*-infected RAW 264.7 macrophages. (A) Knockdown efficiency for each SR and hnRNP factor as measured by RT-qPCR. Data is shown as hnRNP/SRSF expression, relative to *Actb*, compared to SCR control cells. Ratios are the mean of 3 biological replicates and error bars show standard deviation. (B) (top) Overlap of differentially expressed genes (DEGs) between uninfected and *Salmonella*-infected RAW 264.7 macrophages (4h post-infection; MOI = 10) via Venn Diagram. (bottom) Heatmaps of DEGs from uninfected macrophages (both up- and down-regulated; p<0.05). The values of the same genes in Salmonella-infected cells are shown below, with “blank spots” indicating DEGs that are not significantly changed in *Salmonella*-infected SR/hnRNP knockdown cell lines. Orange represents genes downregulated in knockdown vs. SCR; purple represents genes upregulated in knockdown vs. SCR (colorbar shown to the right). All DEGs defined based on having a statisical significant fold change relative to SCR; p<0.05. (C) Ingenuity pathway analysis (IPA) of functional cellular pathways enriched for DEGs in uninfected SR and hnRNP knockdown macrophages. Pathways enriched in eight or more knockdown cell lines are shown. Enrichment is expressed as (-log (p-value)). (D) As in (C) but in *Salmonella*-infected macrophages. Pathways enriched in eight or more knockdown cell lines are shown except for “Regulation of eIF4 and p70S6K Signaling,” which is shown because it is the only pathway that is common to any SR/hnRNP knockdown cell line in both uninfected and *Salmonella*-infected conditions.

To induce macrophage innate immune gene expression, we infected each of the RAW 264.7 knockdown cell lines alongside two SCR control cell lines with *Salmonella enterica* serovar Typhimurium at an MOI of 10. *Salmonella* was chosen because it elicits a widely-studied innate immune response wherein TLR4 sensing of *Salmonella* lipopolysaccharide (LPS) triggers two distinct transcription factor regulons—NFκB downstream of the MyD88 adapter protein and IRF3 downstream of the adapter TRIF [30]. Because *Salmonella* infection of RAW 264.7 macrophages has previously been shown to elicit similar gene expression responses as LPS treatment at 4h [31], we will be able to compare our data to earlier studies of LPS-treated cells, while benefiting from studying the dynamics of a more physiological and disease-relevant response. We collected total RNA from uninfected and *Salmonella*-infected macrophages at 4h post-infection and performed bulk RNA sequencing via an Illumina HiSeq 4000 sequencer (150bp; paired-end reads). An average of ∼60.2 million raw sequencing reads were generated from three biological replicates (20 million reads per sample) of each knockdown (both in uninfected and *Salmonella*-infected conditions). Reads were aligned, quantified, and analyzed using CLC Genomics Workbench (Qiagen).

To determine if knockdown of SR and hnRNP proteins affected expression of different transcripts in uninfected vs. *Salmonella*-infected macrophages, we identified transcripts whose expression was significantly altered (p<0.05; up- or down-regulated) in knockdown cell lines relative to controls and generated Venn diagrams to visualize overlap between affected genes in uninfected (UN) and *Salmonella*-infected (+SAL) macrophages. We deemed these “Differentially Expressed Genes” or DEGs. On average, about 1/3 of DEGs were common to both conditions in the majority of SR and hnRNP knockdown macrophage cell lines and knockdown of most factors impacted between 200-400 genes in both uninfected or *Salmonella*-infected macrophages. Knockdown of hnRNP F, which altered the abundance of 1943 genes in uninfected macrophages and 732 genes in *Salmonella*-infected macrophages, was a notable exception (Table S1 contains all gene expression changes p<0.05).

Upon *Salmonella* infection of macrophages, many innate immune transcripts including cytokines, chemokines, and antimicrobial mediators are dramatically upregulated. Because the vast majority of these genes are either expressed at very low levels or not at all in uninfected macrophages, it was not surprising that many of the DEGs in *Salmonella*-infected cells were not represented amongst the uninfected DEGs (Fig. 1B). However, we found it curious that DEGs in uninfected SR/hnRNP knockdown cells, many of which (e.g. *Hpgd, Lpl, Lars2, Lnpep, Fnip1* (Table S1)) are involved in basic cellular homeostasis and metabolism, are not differentially expressed in SR/hnRNP knockdowns compared to SCR controls in the context of *Salmonella* infection (Fig. 1B heatmaps, UN vs. SAL). Previous studies in yeast have demonstrated that sequestration of the spliceosome can play a powerful role in regulating the cellular response to stress, specifically via downregulation of ribosomal biogenesis [32-36]. Therefore, we hypothesized that downregulation of SR/hnRNP-dependent DEGs (either at the level of transcription or mRNA turnover) in uninfected macrophages could be responsible for loss of SR/hnRNP-reliance during *Salmonella* infection. To test this prediction, we manually cross-referenced SR/hnRNP-sensitive genes against genes whose expression was downregulated in *Salmonella*-infected SCR control macrophages (Table S1). 365 genes were downregulated 2-fold or more (p<0.05) at 4h post-*Salmonella* infection. Many of these genes (e.g. *Lhfpl2, Bhlhe41, Hyal1*, and *Tbc1d2*) have previously been reported as differentially expressed in M1 vs. M2 macrophages and their downregulation likely represents M1 polarization that occurs following *Salmonella* infection [37]. Only a handful of these 365 downregulated genes were among the SR/hnRNP-sensitive genes in uninfected macrophages and we confirmed lack of downregulation for several representative DEGs (e.g. *Bnip3, Id2, Hpgd*) in SR/hnRNP knockdowns by RT-qPCR (Fig. S1B-E). In fact, none of our ten knockdown cell lines showed more than 6.3% of uninfected DEGs being downregulated upon *Salmonella* infection (Fig. S1F). Therefore, we can conclude that SR/hnRNP-dependent DEGs in uninfected cells are generally not downregulated upon infection, although their abundance is no longer differentially impacted by SR/hnRNP knockdown compared to SCR controls. This finding begins to suggest that SR proteins and hnRNPs are functionalized such that the nature of their target transcripts changes in the context of macrophage infection.

As another measure of how SR/hnRNPs differentially influence gene expression in uninfected vs. *Salmonella*-infected macrophages, we performed Ingenuity Pathway Analysis (Qiagen) to identify pathways enriched for SR/hnRNP-sensitive DEGs. In uninfected macrophages, we observed significant enrichment for DEGs in pathways related to translation initiation, mTOR signal-ing, and phagosomal maturation (Fig. 1C, and Table S1 for full list), which paralleled previous anal-yses in hnRNP M knockdown macrophages [38]. IPA of the set of 365 genes described above that are downregulated (>2-fold down) upon *Salmonella* infection revealed no overlap between pathways enriched for SR/hnRNP-sensitive DEGs and those enriched for downregulated genes (Fig. S1G), supporting our conclusion that SR/hnRNP target genes (Fig. 1C) are not globally downregulated upon infection.

Major pathways enriched for SR/hnRNP DEGs in *Salmonella*-infected macrophages were generally related to innate immune responses and macrophage activation, namely “Recognition of Bacteria and Viruses by Pattern Recognition Receptors,” “Communication between Innate and Adaptive Immune Cells,” and “Granulocyte Adhesion and Diapedesis” (Fig. 1D). Consistent with our observation that many DEGs identified in uninfected macrophages lose reliance on particular SR/hnRNPs for proper expression upon *Salmonella*-infection, the only pathway that was significantly enriched for DEGs in both conditions was the translation/mTOR related pathway “Regulation of eIF4 p70S6K,” which remained significantly enriched for DEGs in SRSF6 and SRSF9 in *Salmonella*-infected macrophages (Fig. 1D). Altogether, these analyses demonstrate that SR and hnRNP targets differ dramatically between uninfected and *Salmonella*-infected macrophages and suggest that a gene’s reliance on a particular SR or hnRNP can be altered by macrophage activation (even if its abundance is unchanged by *Salmonella*-infection).

### SR and hnRNPs can both repress and activate innate immune gene expression during *Salmonella* infection

Having observed distinct sets of transcripts whose expression were altered by loss of splicing regulatory factors in uninfected and *Salmonella*-infected macrophages, we set out to identify functional trends in these data. To begin, we chose several of the categories enriched for SR/hnRNP-sensitive DEGs from our IPA analysis (Fig. 1C and 1D) and examined how specific DEGs were impacted by knockdown of individual SR/hnRNPs. In uninfected macrophages, there was enrichment of DEGs in the functional category “Regulation of eIF4 and P70S6K (P70-S6 Kinase 1) Signaling,” which is comprised of ribosomal protein genes and related translation factors (Fig. 2A). Knockdown of hnRNPs generally led to upregulation of DEGs in this category (Fig. 2A; red circles and lines) whereas knockdown of SR proteins led to lower steady state abundance (Fig. 2A; grey circles and lines). The majority of “hits” in this category were differentially expressed in both SR and hnRNP knockdown macrophage cell lines, suggesting that the main job of SR/hnRNP splicing regulatory proteins in resting macrophages is controlling the expression of components of the translation machinery. In *Salmonella*-infected macrophages, we only see enrichment of DEGs in this category in SRSF6 and SRSF9 knockdowns, and the overall number of DEGs is much lower, suggesting that reliance on these splicing factors for maintaining proper gene expression levels shifts away from genes related to translational regulation upon macrophage activation (Fig. 2B).

**Figure 2:**
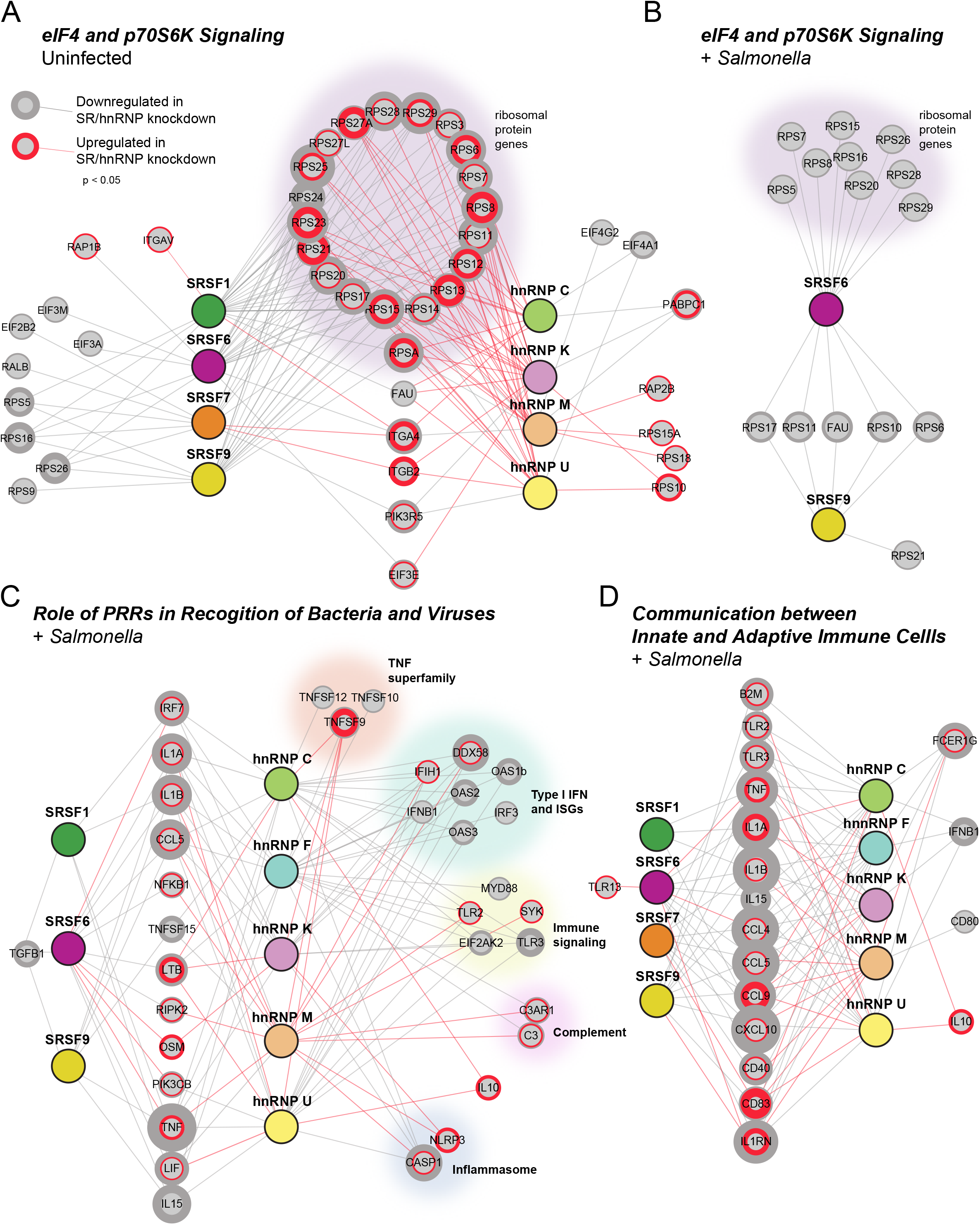
SR/hnRNP-dependent DEGs differ dramatically between uninfected and *Salmonella*-infected macrophages. (A) Network diagrams showing DEGs from the IPA category “eIF4 and p70S6K Signaling” from Figure 1C, showing DEGs from each SR/hnRNP knockdown cell line in uninfected RAW 264.7 macrophages. Only SR/hnRNPs that showed DEG enrichment for the eIF4 pathway are shown. (B) As in (A) but for DEGs measured in *Salmonella*-infected cells. Only SRSF6 and SRSF9 knockdown had DEGs that were statistically enriched in this IPA category during *Salmonella* infection. (C) As in (B) but for DEGs in *Salmonella*-infected SR/hnRNP knockdown cell lines enriched in the IPA category “Role of Pattern Recognition Receptors in Recognition of Bacteria and Viruses.” (D) As in (C) but for the IPA category “Communication between Innate and Adaptive Immune Cells.” Red circles and lines connect SR or hnRNPs with target genes whose expression was upregulated in knockdown vs. SCR control macrophages. Grey circles and lines connect SR or hnRNPs with target genes whose expression was downregulated in knockdown vs. SCR control macrophages. Cut-off for inclusion in the IPA was p<0.05 for differential expression between knockdown and SCR cells.

Looking at IPA categories enriched for DEGs only during *Salmonella* infection (e.g. “Role of Pattern Recognition Receptors in Recognition of Bacteria and Viruses” (Fig. 2C) and “Communication between Innate and Adaptive Immune Cells” (Fig. 2D)), we noticed a similar trend whereby knockdown of hnRNPs upregulates many more genes than SR knockdown. While many innate immune genes in these categories were again, differentially expressed in response to both SR and hnRNP knockdown, there was noticeable enrichment for hnRNP-dependent regulation of factors related to type I IFN expression and ISGs (*Ifih1* (which encodes the viral RNA sensor MDA5), *Ifnb1, Oas1b, Oas2, Oas3, Irf3, Ddx58* (which encodes the viral RNA sensor RIG-I)), several members of the TNF superfamily, and the anti-inflammatory cytokine *Il10*, which is significantly more abundant in both hnRNP C and U knockdowns, relative to control cells, (+4.6 and +3.1-fold change, respectively).

To take a closer look at how knockdown of each SR/hnRNP impacts the macrophage transcriptome during *Salmonella* infection, we quantified the number of transcripts whose expression was up- or down-regulated in the absence of each SR or hnRNP, compared to controls (p<0.05). As visualized in Fig. 3A-E (SRs) and 4A-E (hnRNPs), most splicing factors queried were able to act as both activators and repressors of gene expression in *Salmonella*-infected macrophages, as roughly equal numbers of genes had positive and negative abundance changes in most knockdowns. hnRNP F was a notable exception to this rule, as almost all genes affected by loss of hnRNP F are significantly downregulated (680 genes down, 52 genes up), suggesting an especially important role for hnRNP F in activating innate immune gene expression. Similar trends were observed in uninfected macrophages, whereby roughly equivalent numbers of genes were up and down-regulated in SR/hnRNP knockdown cells (Fig. 1B and Table S1).

**Figure 3:**
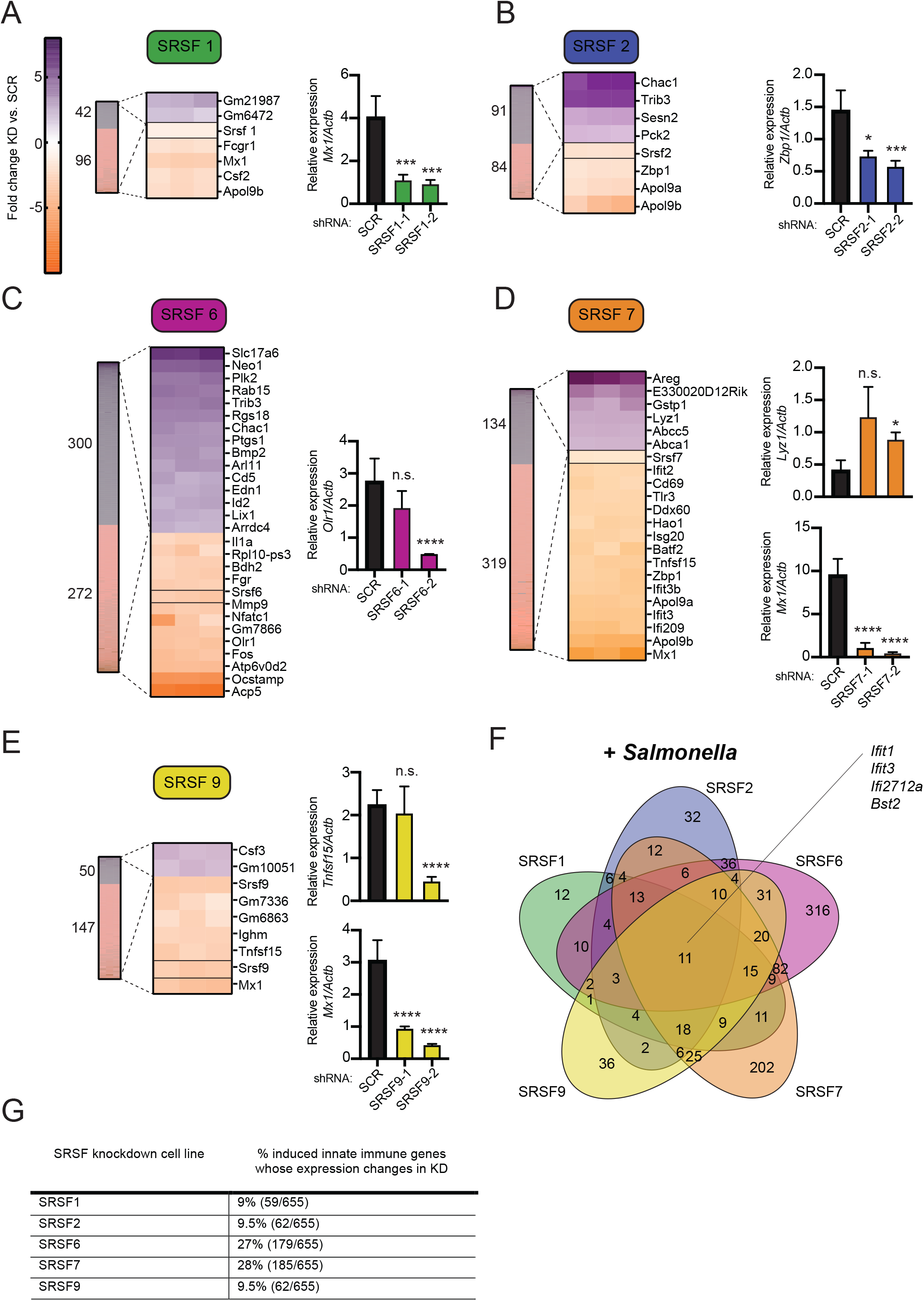
Knockdown of SR family members causes up- and down-regulation of gene expression in *Salmonella*-infected macrophages. (A) On left, heatmap represents all up- and down-regulated genes in SRSF1 knockdown macrophages relative to SCR control at 4h post-*Salmonella* infection (p<0.05). Zoom in represents the top 5% up- and down-regulated genes (fold change). Numbers next to heatmap indicate the number of up- (purple) or down- (orange) regulated genes in SRSF1 knockdown cell lines vs. SCR. On right, RT-qPCR validation of expression of a representative DEG (*Mx1*) in two knockdown cell lines vs. SCR control cells. (B) As in (A) but for SRSF2; RT-qPCR of *Zbp1*. (C) As in (A) but for SRSF6; RT-qPCR of *Olr1*; (D) As in (A) but for SRSF7; RT-qPCR of *Lyz1* and *Mx1*; (E) As in (A) but for SRSF9; RT-qPCR of *Tnfsf15* and *Mx1*. (F) Venn diagram of DEGs common to one or more SR knockdown cell line (p<0.05). Interferon stimulated genes (ISGs) whose expression is impacted by loss of all five SRSF proteins are highlighted. (G) Percentage of all genes induced at 4h post-*Salmonella* infection (>2.0-fold) that are differentially expressed in each SRSF knockdown macrophage cell line (p<0.05). For all RT-qPCRs, values are the mean of 3 biological replicates and error bars indicate standard deviation. * = p<0.05; ** = p<0.01; *** = p<0.005; **** = p<0.001; n.s. = not statistically significant.

To identify the most impacted DEGs, we generated heatmaps that show the top 5% of up- and down-regulated genes (p<0.05) in each *Salmonella*-infected SR/hnRNP knockdown macrophage cell line compared to SCR, with the expression level of each shRNA-targeted SR/hnRNP gene manually included as a reference (Fig. 3A-E and Fig. 4A-E). These heatmaps show clear hyper- or hypo-induction of many critical innate immune genes in SR/hnRNP knockdowns, including genes encoding nitric oxide synthase (*Nos2*) and many interferon stimulated genes (ISGs), including the viral restriction factor *Mx1*, Apolipoprotein 9 (*Apol9a/b*), and members of the IFIT (Interferon-induced protein with tetratricopeptide repeats) family of proteins that interfere with protein synthesis during viral infection (i.e. *Ifit2, Ifit3, Ifit1bl1*). We validated differential expression of top DEGs by RT-qPCR (Fig. 3A-E and Fig. 4A-E) using two different knockdown cell lines for each SR/hnRNP (knockdown efficiency was validated by RT-qPCR and, in cases where antibodies specific for endogenous proteins were available, by immunoblot (SRs in Fig. S2 and hnRNPs in Fig. S3).

**Figure 4:**
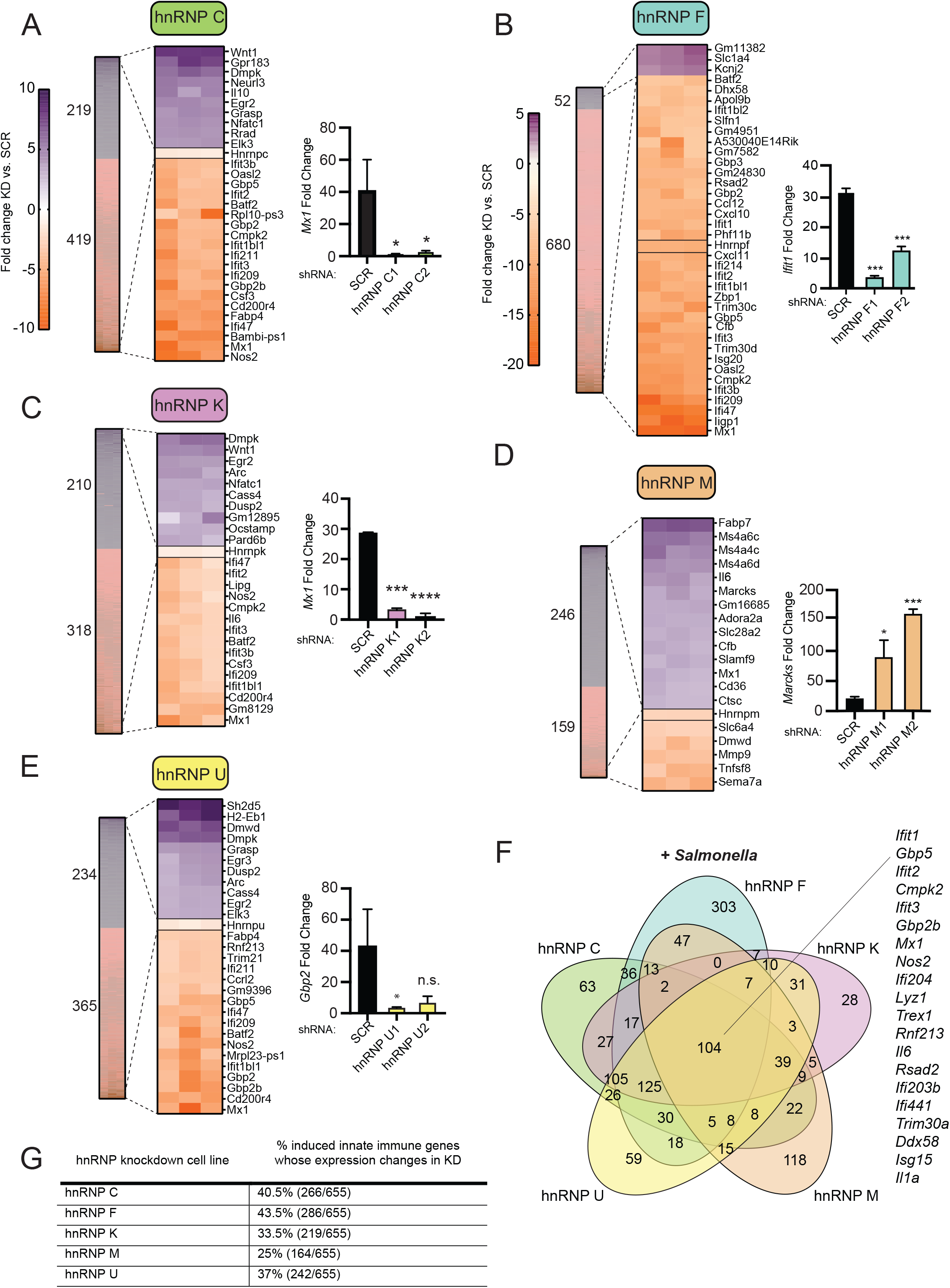
Knockdown of hnRNP family members causes up- and down-regulation of gene expression in *Salmonella*-infected macrophages. (A) On left, heatmap represents all up- and down-regulated genes in hnRNP C knockdown macrophages relative to SCR control at 4h post-*Salmonella* infection (p<0.05). Zoom in represents the top 5% up- and down-regulated genes (fold change). Numbers next to heatmap indicate the number of up- (purple) or down- (orange) regulated genes in SRSF1 knockdown cell lines vs. SCR. On right, RT-qPCR validation of expression of a representative DEG (*Mx1*) in two knockdown cell lines vs. a SCR control. (B) As in (A) but for hnRNP F; RT-qPCR of *Ifit1*. (C) As in (A) but for hnRNP K; RT-qPCR of *Mx1*. (D) As in (A) but for hnRNP M; RT-qPCR of *Marcks*; (E) As in (A) but for hnRNP U; RT-qPCR of *Gbp2*; (F) Venn diagram of DEGs common to one or more hnRNP knockdown cell line (p<0.05). Interferon stimulated genes (ISGs) whose expression is impacted by loss of all five hnRNP proteins are highlighted. (G) Percentage of all genes induced at 4h post-*Salmonella* infection (>2.0-fold) that are differentially expressed in each hnRNP knockdown macrophage cell line (p<0.05). For all RT-qPCRs values are the mean of 3 biological replicates and error bars indicate standard deviation. * = p<0.05; ** = p<0.01; *** = p<0.005; n.s. = not statistically significant.

We next asked whether certain DEGs were shared targets of more than one SR or hnRNP, as a way to provide evidence for cooperativity between factors and to identify families of genes particularly sensitive to loss of these factors. Interestingly, we observed more shared DEGs amongst hnRNP (104 DEGs common to all five knockdowns) vs. SRSF knockdown cell lines (11 DEGs common to all five knockdowns), with another 125 DEGs shared between hnRNP C, K, and U, in particular (Fig. 3F and 4F). This result echoed previous global analyses of hnRNP A1, A2/B1, F, H1, M, and U targets in human 293T cells, which described considerable cooperation between hnRNP family members [39]. Many of the DEGs shared between multiple SR knockdown cell lines and multiple hnRNP knockdown cell lines are characterized as ISGs, as defined by [40-42] (Fig. 3F and 4F), suggesting that genes in this family, which play key roles in regulating antiviral responses and are expressed by members of the IRF and STAT families of transcription factors, are particularly reliant on SRs/hnRNPs to regulate their induction following inflammatory triggers. Overall, we found that between 25-43% of all genes that are induced >2-fold at 4h post*Salmonella* infection had altered gene expression in one or more hnRNP knockdown cell line and 9-25% in one or more SR knockdown cell line (Fig. 3G and 4G). These data demonstrate that the innate immune response relies heavily on splicing regulatory proteins to control expression/abundance of induced genes (in particular, ISGs) and suggest a privileged role for hnRNP proteins in this response.

### SR/hnRNP-mediated alternative splicing events are not common in DEGs

Data presented so far generally argue against a global up- or down-regulation of pre-mRNA splicing in *Salmonella*-infected macrophages and instead support a model whereby individual SRs and hnRNPs dictate RNA processing decisions for particular transcripts. Both SR and hnRNP proteins have been implicated in many steps of gene expression and RNA processing, from chromatin remodeling and transcription, to constitutive and alternative splicing, to mRNA export and stability [43-46]. One obvious way in which all of these proteins can shape the steady state transcriptome, and likely innate immune outcomes, is via alternative splicing. Recent work from the Baltimore lab has shown that innate immune transcripts are frequently regulated by alternative splicing events that introduce so-called “poison exons” that introduce premature stop codons and target transcripts to nonsense mediated decay [21]. To determine whether alternative splicing events could explain differential gene expression in SR/hnRNP knockdown macrophages, we employed an algorithm to identify and quantify local splicing variations (LSV) called Modeling Alternative Junction Inclusion Quantification (MAJIQ) [47]. MAJIQ allows identification, quantification, and visualization of diverse LSVs, including alternative 5′ or 3′ splice site usage, exon skipping, and intron retention across different experimental conditions.

We used MAJIQ to quantify LSVs between SR/hnRNP knockdown and SCR control cell lines in both uninfected and *Salmonella*-infected conditions, generating a large dataset of SR/hnRNP-dependent alternative splicing changes. MAJIQ identified hundreds of statistically significant alternative splicing changes (probability [∣delta PSI∣, ≥10%] >95%) in all SR and hnRNP knockdown macrophages, both in uninfected cells and during *Salmonella* infection. We observed all types of alternative splicing changes, with the majority of changes being categorized as exon skipping events, consistent with the canonical roles of SR and hnRNPs in enhancing or repressing exon inclusion, respectively (Fig. 5A) [48-50]. There were no dramatic differences between the overall number of LSVs between different SR/hnRNPs nor major differences in the number of LSVs in uninfected vs. *Salmonella*-infected macrophages (Fig. 5A). One exception to this trend was that loss of hnRNP F (Fig. 5A, teal) impacted more LSVs of all varieties (intron retention, exon skipping, alternative 3’SS, alternative 5’SS) in uninfected vs. *Salmonella*-infected cells. In uninfected cells, loss of hnRNP F impacted 2807 exon skipping events (vs. 835 in *Salmonella*-infected), 803 intron retention events (vs. 219), 597 alternative 3’SS events, (vs. 177) and 641 alternative 5’SS events (vs. 201) compared to 835, supporting our hypothesis that macrophage activation and differential phosphorylation can alter the function of regulatory splicing factors, perhaps by changing their protein binding partners, RNA targets, or subcellular localization. Generally, we observed only a fraction of DIU events were common to uninfected and infected macrophages (Fig. S4B), which paralleled what we saw for DEGs between the two conditions.

**Figure 5:**
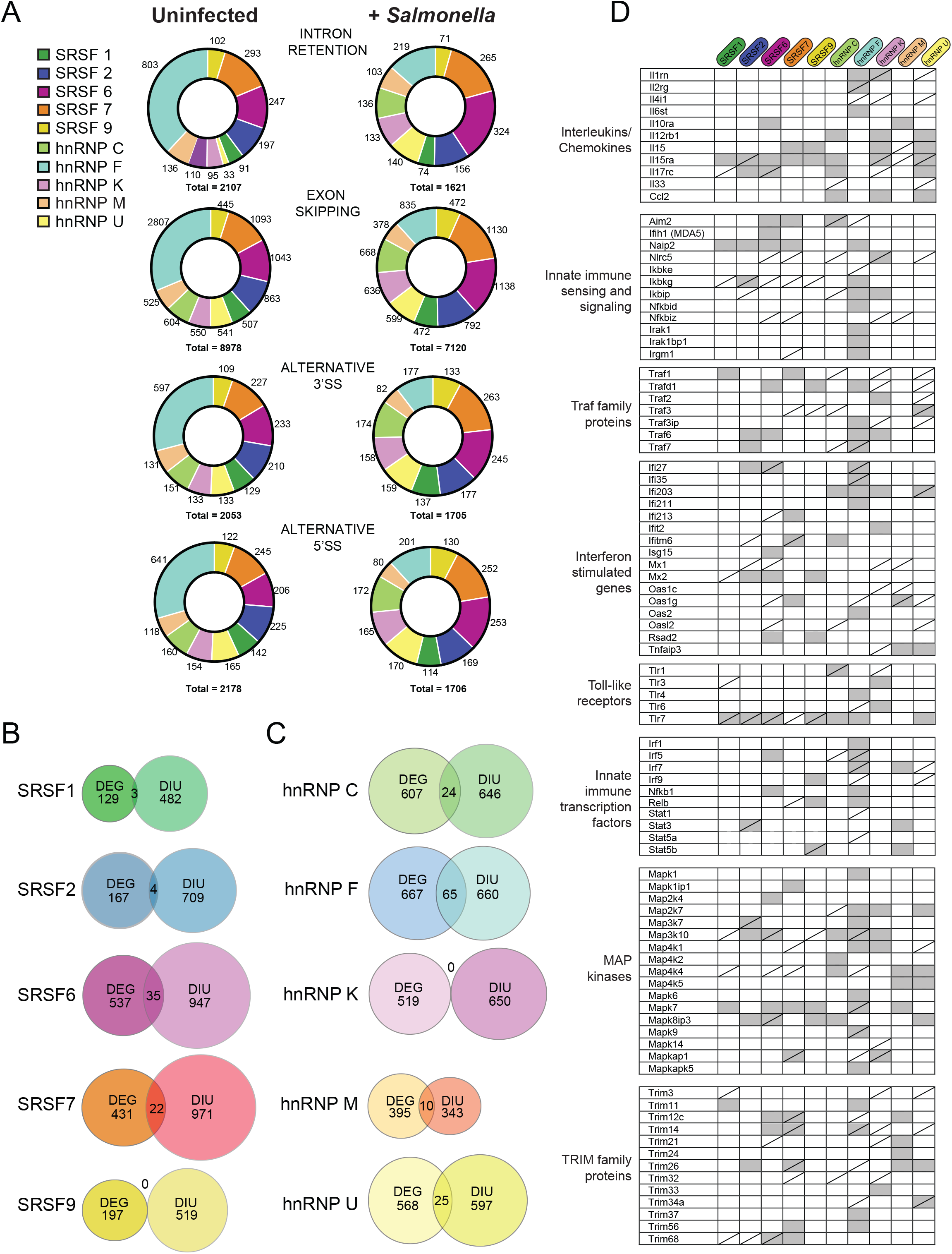
Local splicing variations are abundant in SR/hnRNP knockdown macrophages, but they do not preferentially occur in SR/hnRNP-dependent differentially expressed genes. (A) Quantitation of intron retention, exon skipping, alternative 3’ and 5’ splice site events in uninfected and *Salmonella*-infected SR and hnRNP knockdown macrophages (probability [∣delta PSI∣, ≥10%], >95%). (B) Overlap between DEGs (p<0.05) and genes with differential isoform usage (DIU) as defined by MAJIQ LSVs (PSI ≥10%) in each SRSF knockdown macrophage cell line +*Salmonella* infection. (C) As in (B) but for hnRNP knockdown cell lines. (D) Identification of innate immune-related genes with DIU in each SR/hnRNP knockdown macrophage cell line vs. SCR control. Genes included in chart have one or more significant LSV by MAJIQ analysis. Shaded boxes indicate an LSV occurs preferentially in uninfected knockdown macrophages; slashes indicate an LSV occurs preferentially in *Salmonella*-infected macrophages. Genes were manually annotated into different innate immune categories.

To investigate a role for alternative splicing in regulating innate immune gene expression in *Salmonella*-infected macrophages, we calculated the overlap between DEGs and genes with one or more LSV (which we will refer to as differential isoform usage or DIU). Surprisingly, we saw almost no overlap between genes whose expression was changed in SR/hnRNP knockdowns vs. those whose alternative splicing was changed (DEG vs. DIU). This was the case for both uninfected (Fig. S4A) and *Salmonella*-infected conditions (Fig. 5C and 5D). Likewise, we observed little to no enrichment for SR/hnRNP-dependent alternative splicing events in genes related to innate immune pathways via IPA. In fact, no pathway was enriched for MAJIQ hits more than -log(p-value) = 5 (Tables S2 (SR) and S3 (hnRNP)), suggesting that SR/hnRNP-dependent alternative splicing changes in macrophages do not occur in functionally related genes from any particular pathway. Other studies also found a lack of overlap between steady state transcript levels and alternative splicing events in *Salmonella*-infected human monocytes [20] and influenza-infected A549 cells [51]. Together, these data suggest that changes to constitutive intron removal and/or alternative splicing of an upstream regulator that influences transcription or stability of DEGs are more likely explanations for how loss of SR/hnRNPs influences steady state RNA expression levels in cells.

To address the latter possibility that innate immune gene expression is globally impacted by an alternatively spliced isoform of a global regulator (innate immune sensor or signaling factor), we manually cataloged genes with one or more instance of DIU in SR/hnRNP knockdown cell lines into categories related to immune function and asked whether SR/hnRNP knockdown led to distinct instances of DIU in uninfected and/or *Salmonella*-infected macrophages (Fig. 5D). While differential expression of innate immune genes was generally unique to *Salmonella*-infected mac-rophages, SR/hnRNP-dependent alternative splicing changes of immune genes were detected in each of the experimental conditions queried. Some genes cells (e.g. *Naip2, Traf6, Il12rb1, Nfkb1*, and *Trim56*) showed alternative splicing changes only in uninfected, others (e.g. *Tlr7, Il15ra, Trim14*) were alternatively spliced in both conditions, and others (e.g. *Trim3, Mx1, Traf3, Ikbkg*) were only impacted by loss of SR/ hnRNPs in the context of *Salmonella* infection. While the SR/hnRNPs queried certainly contribute to alternative splicing of innate immune signaling mole-cules and regulatory factors, we can conclude that it is unlikely that the generation of alternatively spliced isoforms is the main driver of transcript abundance changes in SR/hnRNP knockdown macrophage cell lines.

### Select innate immune genes are reliant on SR/hRNP proteins for maintaining proper expression levels

While our transcriptomics data suggest that each of the SR/hnRNPs queried impacts its own cohort of transcripts, we noticed that several innate immune genes were “hit” in multiple SR/hRNP knockdown cell lines, including *Il1a* (a pro-inflammatory cytokine that produces numerous inflammatory signaling events through the IL1 receptor [52]), *Nos2* (nitric oxide synthase, an important antimicrobial effector molecule [53]), and *Mx1* (a viral restriction factor [54]). To begin to understand why the abundance of these genes relies heavily on SR/hnRNPs, we first validated hyper- and hypo-induction of these genes by RT-qPCR in *Salmonella*-infected samples from all ten of our knockdown macrophage cell lines and our two SCR controls. We observed that *Il1a* induction was higher in the absence of hnRNP C, hnRNP K, SRSF7, and to limited extent, hnRNP U (Fig. 6A). Likewise, *Nos2* expression varied dramatically in different SR/hnRNP knockdown macrophages, with hnRNP C, K, U and SRSF1 being required for maximal induction of *Nos2* and factors like hnRNP M, SRSF6 and SRSF7 contributing to *Nos2* repression (Fig. 6B). Curiously, upregulation of *Nos2* in SRSF6 knockdown cells was seen both at rest (Table S1) and following *Salmonella* infection (Fig. 6B), suggesting that loss of SRSF6 somehow stimulates basal innate immune activation. *Mx1* expression was lower in all of the hnRNP as well as in SRSF1 and 7 knockdowns compared to scramble controls, except for hnRNP M (Fig. 6C), consistent with our previously characterized role for this factor in repressing *Mx1* expression [38]. Having confirmed reliance of each of these genes on particular SR/hnRNPs 4h post-Salmonella infection, we turned to a computational prediction method called RBPFinder to unbiasedly identify potential binding sites for each of our SR and hnRNPs of interest [55]. RBPFinder detected hundreds of potential binding sites for these RNA binding proteins in *Il1a, Nos2, Mx1, Il10* (Fig. 6D-E), and *Ddx58* (Fig. 6F-G). For each case we examined, if a transcript’s abundance was impacted by loss of a particular SR/hnRNP, it encoded one or more binding sites for that factor (*Ila1*: hnRNP C, K, U, SRSF7; *Nos2*: hnRNP C, K, M, U, SRSF1, 6, 7; *Mx1*: hnRNP C, F, K, M, U, SRSF1, 7, 9; *Il10*: hnRNP C and U; *Ddx58*: hnRNP C, F, K, M, U, and SRSF7). However, the presence of a binding site was not sufficient to predict reliance on a splicing factor for regulation of a transcript’s induction: for example, our analysis did not identify major changes in *Il1a* expression in the absence of SRSF1, 2, or 9, even though *Il1a* encodes predicted SRSF1, 2, and 9 binding sites (Fig. 6A). While this analysis on its own is merely correlative, it does begin to suggest that exonic and intronic splicing enhancers/silencers are enriched in innate immune transcripts that consequently rely on SR/hnRNPs for regulation.

**Figure 6:**
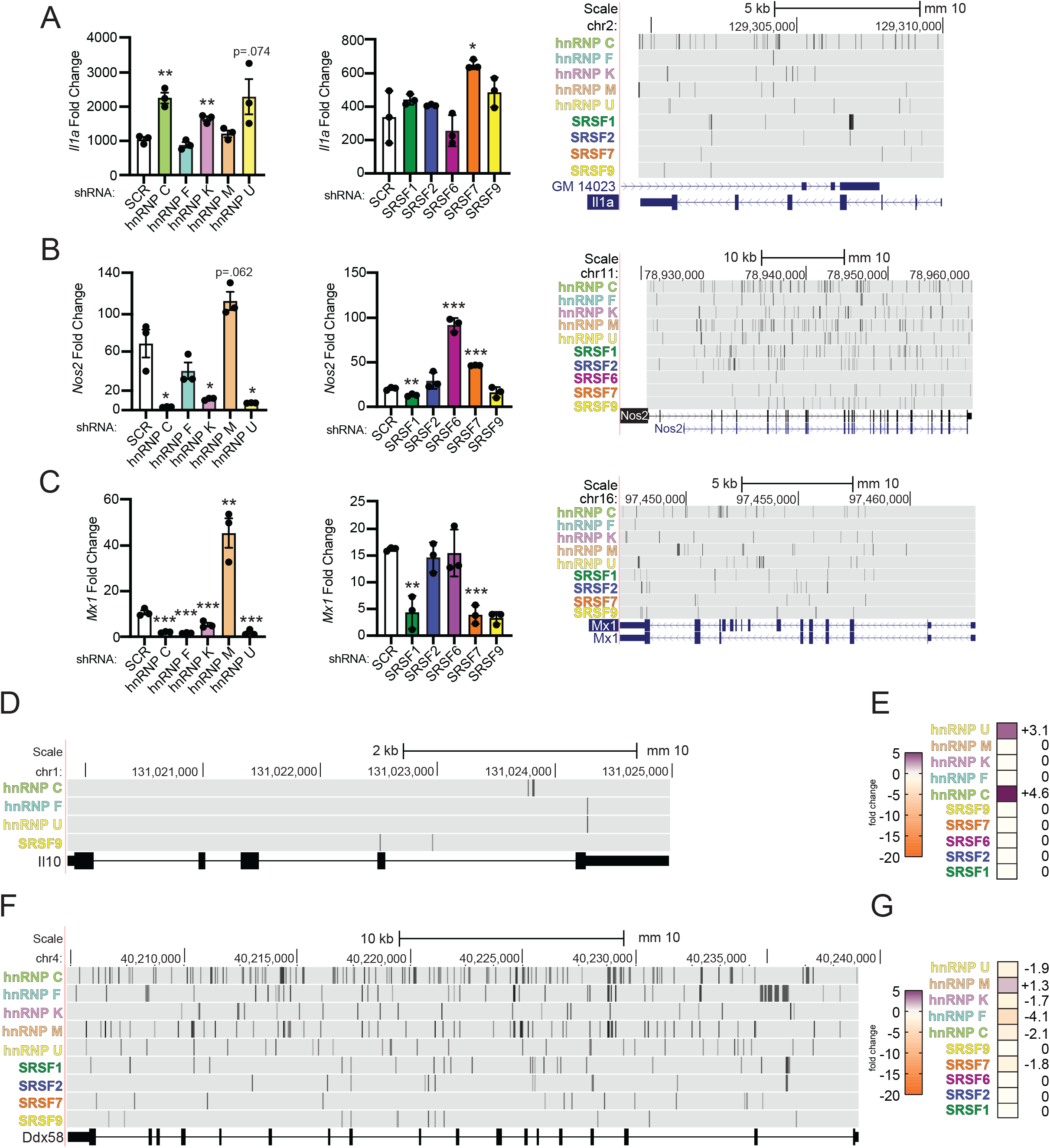
Innate immune genes vary in their reliance of SR/hnRNP family members for proper induction in *Salmonella*-infected RAW 264.7 macrophages. (A) (left) RT-qPCR of *Il1a* abundance relative to *Actb* in *Salmonella*-infected SRSF and hnRNP knockdown macrophages (shown as fold change relative to uninfected). (right) Gene diagram of Il1a from UCSC genome browser with RBPFinder tracks for SRSF and hnRNP predicted binding sites. An SRSF/hnRNP track was included if the factor had one or more predicted binding sites in the gene queried. Mouse genome (GRCm38/mm10); high stringency; conservation filter applied). (B) As in (A) but for *Nos2*. (C) As in (A) but for *Mx1*. (D) Gene diagram of Il10 from UCSC genome browser with RBPFinder tracks for SRSF and hnRNP predicted binding sites. (E) Differential gene expression of *Il10* in each SRSF/hnRNP knockdown cell line relative to SCR. Zeros indicate that there was not a statistically significant difference in *Il10* expression in the knockdown cell line vs. SCR. (F) As in (D) but for Ddx58. (G) As in (E) but for *Ddx58*. For all RT-qPCRs, values are the mean of 3 biological replicates and error bars indicate standard deviation. * = p<0.05; ** = p<0.01; *** = p<0.005.

### A gene’s induction level, length, and number of exons/introns do not correlate with a transcript’s reliance on SR/hnRNPs for proper induction during *Salmonella* infection

Having observed a lack of correlation between DEGs and DIU in each splicing factor knockdown, we were eager to see if we could identify anything common to SR/hnRNP-sensitive innate immune genes. We hypothesized that genes whose expression is the most upregulated in response to *Salmonella* infection could be more sensitive to loss of SR/hnRNPs, perhaps via a need to sequester potentially rate-limiting spliceosome components. To address this possibility, we ranked all genes induced in *Salmonella*-infected SCR control macrophages and looked to see how each of these genes was expressed in our SR/hnRNP knockdown cell lines (Fig. 7A). Consistent with previous reports, we observed dramatic upregulation of hundreds of macrophages genes 4h post-*Salmonella* infection, with some genes like *Il1a* and *Il1b* being upregulated approximately 1500-fold. We observed no clear correlation between level of induction/expression level and whether or not a transcript was differentially regulated by loss of an SR/hnRNP. For example, DEGs induced 1000-fold in control cells were impacted by the loss of a similar number of SR/hnRNPs and to a similar magnitude as DEGs induced 5-fold (Fig. 7A; top vs. bottom of heatmap). To examine if other attributes of a gene influenced whether its expression was altered by loss of an SR/hnRNP, we conducted Pearson’s correlation tests to determine the relationship between differential expression (p<0.05) and gene length (Fig. 7B), exon length (total exonic sequence, or sum of all exon nucleotides) (Fig. 7C), intron length (total intronic sequence, or sum of all intron nucleotides) (Fig. 7D), and number of exons (Fig. 7E). We observed little to no correlation between any of these gene attributes and the degree to which a gene’s expression was altered in the hnRNP and SR knockdown cell lines, with all tests generating Pearson correlation coefficients close to zero. Together, these analyses support a model whereby certain innate immune genes rely more on SR/hnRNPs than others by virtue of transcript intrinsic features like enrichment of SR and hnRNP binding sites.

**Figure 7:**
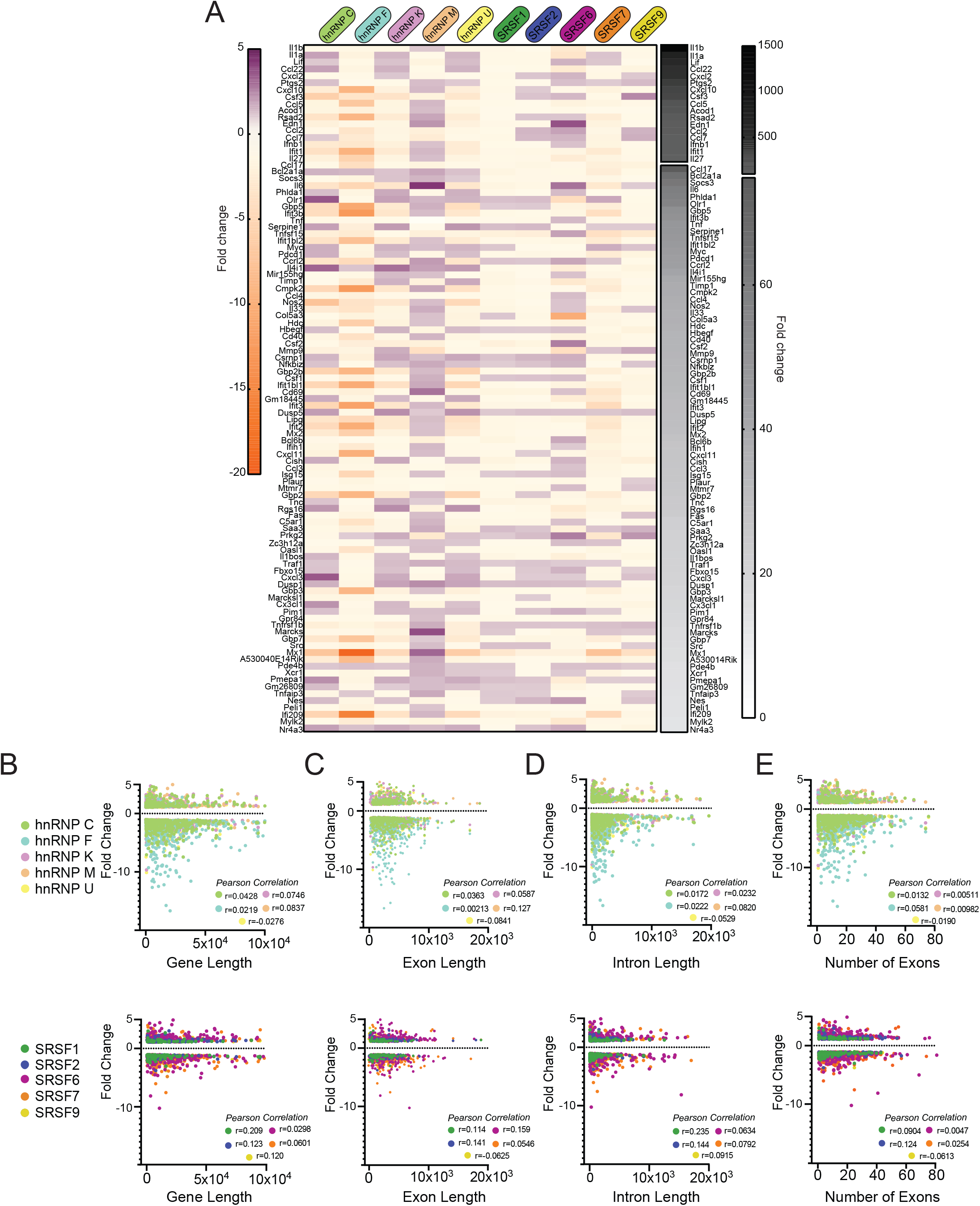
Level of induction, gene length, and number of introns/exons do not positively correlate with a gene’s reliance of SR/hnRNPs for proper expression. (A) (right, black and white) Heatmap of the top 100 genes induced at 4h post-*Salmonella* infection in SCR control RAW 264.7 macrophages. Data shown as fold change SCR +*Salmonella* vs. SCR uninfected. (left) Heatmap of up- or down-regulation (fold change) of each induced gene conferred by hnRNP or SRSF knockdown. Purple, genes upregulated in SR/hnRNP knockdowns; Orange, genes downregulated in SR/hnRNP knockdowns. (B) Scatter plot depicting correlation of the fold change of each DEG vs. coding sequence (CDS) length in *Salmonella*-infected hnRNP (top) and SRSF (bottom) knockdown macrophages. (C) as in (B) but comparison of DEG fold change vs. exon length. (D) As in (B) but comparison of DEG fold change vs. intron length. (E) As in (B) but comparison of DEG fold change vs. number of exons in a DEG. Genes included in analysis were differentially expressed (p<0.05) in each knockdown compared to SCR controls. Y-axes were made all the same to facilitate comparison between multiple knockdown cell lines.

### Rapidly induced innate immune genes are subject to repression from SR and hnRNPs

With no apparent correlation between various gene architecture attributes and SR/hnRNP reliance, we looked to see if we could correlate a gene’s reliance on SR/hnRNP for proper induction and the dynamics of its transcriptional activation as previously described by other studies [6, 7, 10]. One commonly used categorization of innate immune genes is by primary and secondary response genes. Primary response genes are induced rapidly via activation/translocation of existing transcription factors while secondary response genes are expressed with slower kinetics and generally require new protein synthesis of transcription factors before they can be fully induced. Of the 53 LPS-driven primary response genes annotated by [10] (Table S4), 35 of them were up-regulated/more abundant in the absence of one or more SR/hnRNP, suggesting a role for factors like hnRNP C, K, M, U and SRSF6 in repressing the expression of primary response genes (Fig. 8A, top heatmap). A repressive role for the same factors was less evident for secondary response genes, which were mostly downregulated in the absence of the SR/ hnRNPs (except for hnRNP M and SRSF6) (Fig. 8A, bottom heatmap). This analysis begins to suggest that primary response genes may be more reliant on pre-mRNA splicing to control the proper timing and magnitude of their induction than secondary response genes, which rely on multiple other layers of regulation.

**Figure 8:**
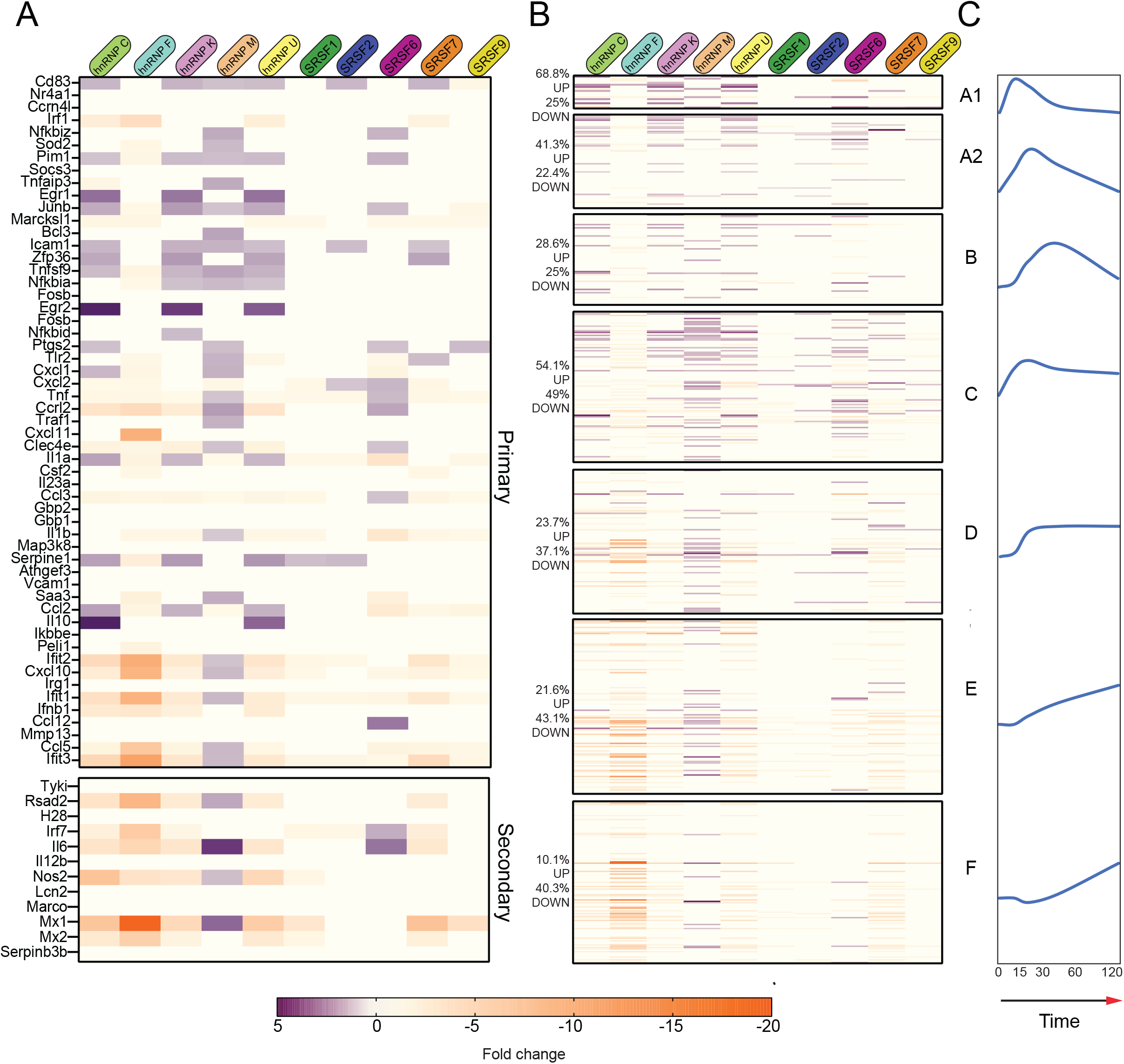
Primary response and early induced innate immune genes are mostly repressed by SR/hnRNPs. (A) Fold change of SR/hnRNP DEGs in *Salmonella*-infected RAW 264.7 macrophages compared to SCR controls with genes categorized as primary and secondary response genes according to Ramirez-Carrozzi et al. 2009. (B) Fold change of SR/hnRNP DEGs in *Salmonella*-infected RAW 264.7 macrophages compared to SCR controls with Lipid A-induced genes categorized on basis of induction kinetics as defined by Bhatt et al., 2012. (C) Schematic representation of induction kinetics of each Group (A1, A2, B, C, D, E, and F) over a 120-minute time course following Lipid A treatment. Adapted from Bhatt et al., 2012.

We next looked at macrophage gene categories as defined by Bhatt et al. [6], which divided RefSeq genes exceeding 400bp in length into 12 clusters based on their pattern of transcript levels in three cellular compartments (chromatin, nucleoplasm, cytoplasm) in primary macro-phages over a time-course of Lipid A treatment. The vast majority of genes we identified as SR/hnRNP-sensitive in *Salmonella*-infected RAW 264.7 macrophages fell into Groups 1-3 (Fig. S5A and Table S5), which are comprised mostly of Lipid A-induced genes. As was the case with the primary response genes, most of the Group 1 genes showed increased expression in SR/hnRNP knock-downs cell lines. We then looked at another categorization by Bhatt et al. [6], that specifically grouped induced genes based on their expression pattern following treatment with Lipid A (Table S6). Genes in Groups A1, A2, and B were characterized by a “burst” of induction, with maximal chromatin-associated reads measured at 15 minutes post-Lipid A for Group A1 genes and slightly slower kinetics measured for Groups A2 and B, respectively. The general dynamics of each of the Groups’ chromatin-reads, based on data in [6] is represented schematically in Fig. 8C. Almost all the genes in Group A1 were differentially expressed in the absence of one or more SR/hnRNP (68.8% genes were upregulated by knockdown of one or more SR/ hnRNP and 25% were down-regulated by one or more knockdown)(Fig. 8B). On average, SR/hnRNP knockdown impacted about 40% of genes in Groups A2-F, with genes in Groups A1, A2, and C tending to increase in abundance in the absence of an SR/hnRNP, and genes in groups D, E, and F, tending to decrease in abundance. Overall, our data argue that hnRNP C, K, M, U, SRSF2, 6, and 7 in particular are important for repressing rapidly induced innate immune genes in Groups A1, A2, B, and C (Fig. 8B).

Because earlier work associated high CpG island-containing promoters with rapid, transi-ently induced primary response genes and low CpG promoters with secondary response genes that display delayed/sustained transcription kinetics, we looked to see if there was any correlation between our SR/hnRNP DEGs and CpG promoter architecture. Generally, DEGs in SR/hnRNP knockdown *Salmonella*-infected macrophages were about evenly split between high CpG and low CpG promoters (Fig. S5B), although Group C DEGs were particularly enriched for CpG promoters (75%) and Group F were notably depleted (23%). Altogether, these data suggest the presence of CpG islands does not dictate whether an innate immune gene relies on SR/hnRNPs for proper induction, although they do support a general role for these regulatory factors in limiting the abundance of primary response genes.

### hnRNP C and hnRNP K share target genes and impact bacterial and viral replication in a similar fashion

Lastly, we wanted to test whether we could use DEG and/or DIU profiles of SR/hnRNP knockdown macrophages to predict if a cell line would be better or worse at controlling infection with a pathogen. We began by applying a simple hierarchical clustering algorithm to calculate similarities in DEG profiles between knockdowns (Cluster 3.0). We found significant similarity between genes affected by loss of hnRNP K and hnRNP U (and to a lesser extent, hnRNP C) in uninfected cells (correlation between K/U: 0.78; correlation between C/K/U: 0.69), as well as in *Salmonella*-infected cells (correlation between K/U: 0.79; correlation between C/K/U: 0.76). Previous studies have shown that hnRNP K binds strongly to poly C stretches of RNA [56, 57] and hnRNP U preferentially binds CUGUGGAU and UGUAUUG motifs [39]. At the amino acid level, hnRNP K and U proteins are only 31% similar in mice (EMBOSS Stretcher Pairwise Sequence Alignment). While their consensus binding motifs argue against their recognizing overlapping sequences, there is evidence from high-throughput studies in humans and mice that hnRNP K and U proteins immunopurify [58] and cofractionate [59, 60] together.

To begin to understand how hnRNP K and U knockdown RAW 264.7 cell lines may be phenotypically similar, we identified several clusters of up- and down-regulated genes common to both cell lines. Two clusters of upregulated genes are highlighted in Figure 9A. Interestingly, Cluster 1 contains mostly ISGs (*Ifi202b, Bst2, Irf7, Ifitm3, Isg15, Ifi44l, Oasl1*) while Cluster 2 is enriched for a diverse group of kinases (*Dmpk, Ripk3*), regulators of GTPase activity (*Gng10, Rgs16, Fgd2*), and mitochondrial related factors (*Pmaip1, Ucp2*). Differential expression of ISGs in uninfected hnRNP K and U knockdown cell lines is notable because it suggests that loss of these factors somehow activates macrophages to upregulate antiviral gene expression. Upon *Salmonella* infection, several of these Cluster 1 ISGs actually become less abundant relative to SCR controls (*Oasl2, Ddx58, Isg15, Usp18*; Cluster 3), again highlighting dysregulation of antiviral genes in the absence of hnRNP K and U. Conversely, Cluster 2 DEGs were more abundant in both uninfected and *Salmonella*-infected hnRNP K and U knockdown macrophages (see Clusters 4 and 5). Overlap between hnRNP K and U DEGs and DIUs is further clarified by Venn Diagrams that show 1/3 shared DEGs and 3/4 shared DIUs.

**Figure 9:**
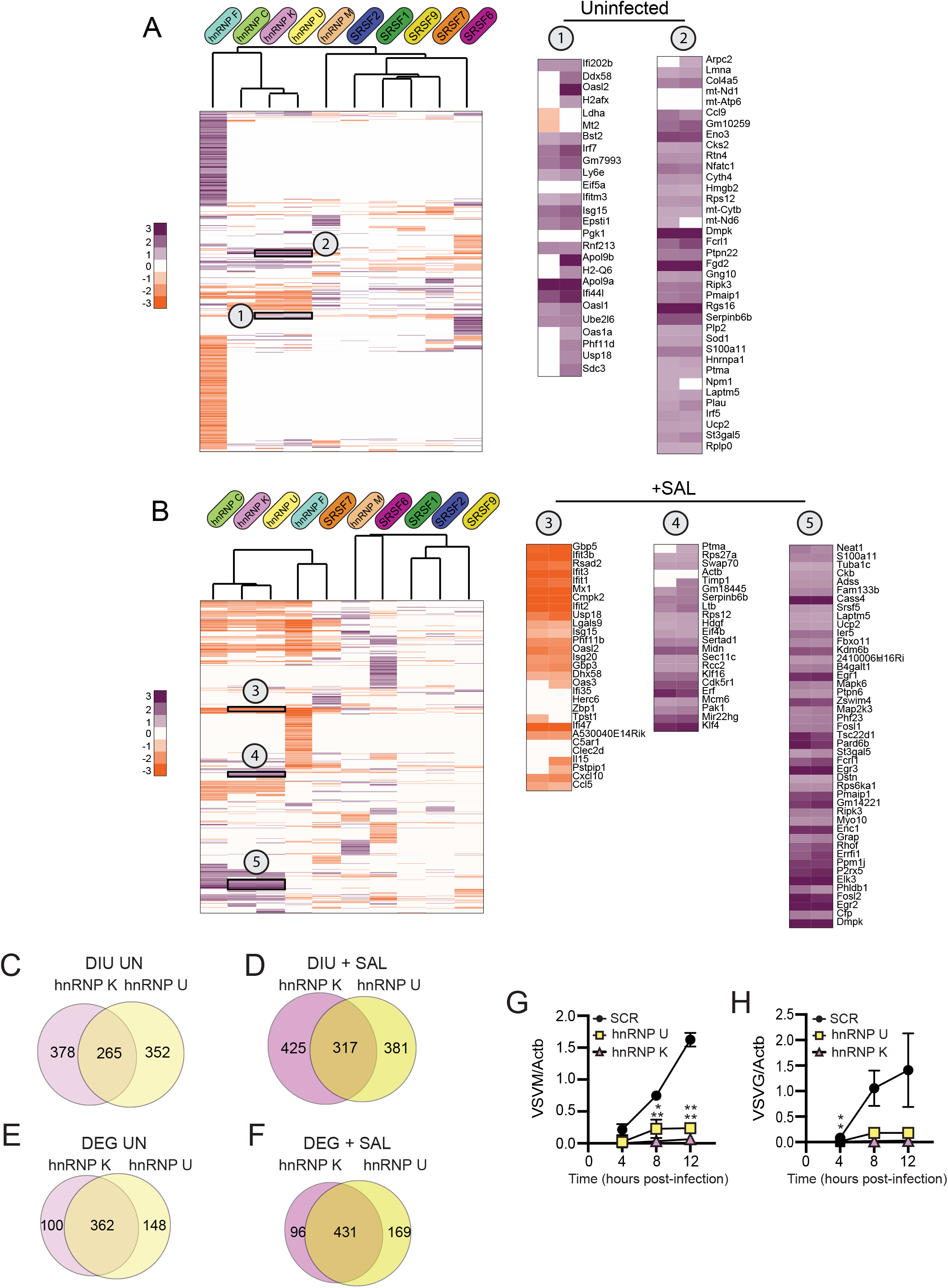
Loss of hnRNP K and hnRNP U impacts similar DEGs and DIU and enhances the ability of RAW 264.7 macrophages to control viral replication. (A) Hierarchical clustering of up- and down-regulated DEGs in uninfected macrophages (SR/hnRNP knockdown vs. SCR). Zoom-ins of clusters of interest (1 and 2) are shown to the right. Correlation of Cluster 1 is 0.84; Cluster 2 is 0.76. To allow for better visualization of DEGs, the scale is set at -3 to +3. (B) As in (A) but in *Salmonella*-infected macrophages. Correlation of Cluster 3 is 0.75; Cluster 4 is 0.78; Cluster 5 is 0.72. (C) Overlap of genes with DIU in uninfected hnRNP K and U knockdown macrophages. (D) As in (C) but comparing DIUs in *Salmonella*-infected macrophages. (E) Overlap of DEGs in uninfected hnRNP K and U knockdown macrophages. (F) As in (E) but comparing DEGs in *Salmonella*-infected macrophages. (G) Viral replication (as measured by RT-qPCR of the VSVM gene) in hnRNP K and hnRNP U knockdown macrophages compared to SCR controls infected with VSV (MOI = 1) at 4, 8, and 12h post-infection. (H) As in (G) but for VSVG. For all RT-qPCRs, values are the mean of 3 biological replicates and error bars indicate standard deviation. * = p<0.05; ** = p<0.01.

Having discovered that loss of hnRNP K and U impacted antiviral gene expression in both uninfected and *Salmonella*-infected macrophages, we asked whether viral replication was impacted at early infection time points in hnRNP K and U knockdown cell lines compared to SCR controls. We infected SCR, hnRNP K and hnRNP U knockdown RAW 264.7 macrophages with VSV at a MOI of 1. VSV is enveloped RNA virus that can replicate and elicit robust gene expression changes in RAW 264.7 cells [61]. Viral replication was measured over a 12h time course by following expression of two viral genes, VSV-G and VSV-M by RT-qPCR. Remarkably, we observed almost no replication of VSV in either hnRNP knockdown cell line at any time point. A similar hyper-restriction phenotype was recently reported for hnRNP K knockdown A549 cell lines infected with influenza virus by Thompson et al. [51]. These results argue strongly for splicing factors contributing to innate immune and infection outcomes in important ways and suggest that some of these factors may work in a cooperative fashion.

## DISCUSSION

While the involvement of splicing factors in regulating steady state gene expression in macrophages should in some ways be a foregone conclusion, our discoveries regarding the diversity of genes impacted by individual splicing factors and the dramatic reliance of some innate immune genes versus others on SR/hnRNPs for establishing or maintaining proper steady state levels, are surprising. Indeed, the fact that over 70% of genes induced 2-fold or more 4h post-*Salmonella* infection are hyper- or hypo-induced in the absence of one or more of the SR/hnRNPs we investigated, highlights a critical role for splicing regulatory proteins in modulating the kinetics and magnitude of gene induction and demands a rethinking of how the innate immune response is post-transcriptionally regulated.

One lingering question raised by these studies relates to the mechanism(s) through which genes are up or down regulated in SR/hnRNP knockdown RAW 264.7 macrophages. While our analysis did identify LSVs in transcripts encoding several transcription factors in the IRF, STAT, and NFκB families (*Irf1, Irf5, Irf7, Irf9*, and *Stat1*), our data does not generally support a model whereby loss of a particular SR/hnRNP results in mis-splicing or functional alteration of a master regulator of a shared transcriptional regulon, apart from hnRNP F and SRSF6, which both appear to dysregulate one or more factors related to interferon signaling or ISG transcription. Likewise, although individual cases likely exist, our data does not support a role for SR/hnRNPs in globally regulating innate immune transcript abundance via differential inclusion of poison exons (as DIU was not enriched in DEGs). Indeed, the most overlap between DIU and DEG we observed for hnRNP F and was still only 10% (Fig. 5)). Thus, the question remains: how do individual SR/hnRNPs activate and/or repress induction of innate immune genes? Identifying the direct RNA binding targets of these splicing factors in uninfected and *Salmonella*-infected macrophages will certainly help answer this question as will defining the subcellular localization of these RNA-binding proteins in the two conditions.

Previous work from our lab, which found hnRNP knockdown leads to hyper-induction of a number of innate immune transcripts following inflammatory triggers, suggests that slowing/inhibiting pre-mRNA maturation may be involved. Specifically, we found that overexpression of hnRNP M promotes accumulation of intron-containing *Il6* pre-mRNAs, while loss of hnRNP M increases removal of *Il6* intron 3 [38]. From these findings we concluded that by repressing constitutive intron removal in *Il6*, hnRNP M acts as a hand break to slow early or spurious activation of this pro-inflammatory cytokine. It is possible that other SR and hnRNPs work in the same way, by contributing to specific constitutive intron removal events that can fine-tune the kinetics of transcript maturation and influence steady state RNA levels. Such a model would predict increased levels of reads from particular introns in DEGs. Although we do not see evidence for this in our RNA-seq data, this is not particularly surprising given the low abundance of intron-containing premRNAs relative to mRNAs. Indeed, an important caveat of these studies is that they were carried out at a single, rather late, time point following *Salmonella* infection. While we chose this timepoint to maximize mRNA transcript accumulation, we may have inadvertently minimized our ability to detect transient accumulation of unprocessed transcript intermediates. Kinetic transcriptome analysis from the Black and Smale labs demonstrates that for most transcripts, pre-mRNA splicing of innate immune transcripts occurs co-transcriptionally but accumulation of nascent pre-mRNA at the chromatin level is generally not evident at time points following 15-30 minutes, except in select cases with especially long transcripts on which splicing catalysis is delayed [6, 7]. Thus, it is possible that for many of our transcripts of interest, the most important contribution of SR/hnRNP proteins to constitutive and/or alternative splicing could occur during that early transcriptional burst. Indeed, our previous experiments looking at hnRNP M support this type of mechanism whereby loss of hnRNP M depressed *Il6* and *Mx1* expression at early time points (2 and 4h) but by 6 and 8h, hnRNP M was dispensable [38]. Thus, future attempts to elucidate the complexities of post-transcriptional control of inflammatory gene induction will want to broaden their scope to include both early and late time-points following macrophage activation.

At the onset of this study, we hypothesized that because SRSF1, 2, 6, 7, 9 and hnRNP C, F, K, M, and U have been shown to be differentially phosphorylated during bacterial and fungal infection of macrophages [27-29], they would impact distinct gene regulons in uninfected vs. S*almonella*-infected cell lines. Indeed, we found that while genes related to ribosome biogenesis and translational regulation change abundance in the absence of these factors in uninfected macrophages, only about half of these “housekeeping” genes were still differentially expressed in knockdowns compared to SCR controls following *Salmonella* infection. We predicted that this discrepancy could be explained by transcriptional downregulation of these genes, perhaps as means to “free up” splicing factors to promote processing of induced innate immune genes. This type of mechanism would be consistent with findings in yeast, which demonstrated that pre-mRNAs compete for splicing factors [62] and that certain introns are evolutionarily maintained specifically to sequester splicing factors, limit RNA processing, and downregulation gene expression in times of stress [32, 33]. Because we measured almost no correlation between genes whose expression was downregulated in *Salmonella*-infected macrophages and those that have and then lose reliance of SR/hnRNPs, we do not believe that transcriptional repression serves as a means to reassign splicing factors from ribosome biogenesis genes to innate immune genes as part of macrophages activation (at least not at the 4h time point we queried). Instead, we propose that functionalization, likely via post-translational modification (PTMs), specifically promotes processing of innate immune transcripts upon macrophage activation. Several studies have linked environmental changes, including starvation [32, 33, 63], heat shock [64-67], and T cell activation [68] to posttranslational modification of splicing factors. For example, dephosphorylation of SRSF10 by the phosphatase PP1 represses splicing and limits gene expression in HeLa cells following heat shock [19, 69, 70] and arginine methylation of hnRNPA1/B2 triggers its export to the cytoplasm where it activates TBK1/IRF3 signaling following infection with a DNA virus [71]. Very recent work from the Lynch lab showed that hnRNP K is redistributed in the nucleus during influenza infection, becoming enriched in nuclear speckles [51]. Indeed, subcellular redistribution is a common trait of SR/hnRNPs during infection [72, 73] and many viruses themselves require RNA binding proteins for the maintenance and processing of their genomes. It is possible that similar mechanisms are at play during macrophage activation, whereby PTMs render SR/hnRNPs more or less likely to engage with certain protein binding partners and/or RNAs, which in turn regulates the proper timing and induction of innate immune genes relative to other targets.

## ACKNOWLEDGEMENTS

We would like to thank members of the Patrick and Watson labs for their critical reading of and feedback on this manuscript. We would also like to thank members of Phillip West’s lab at TAMU COM for help with the VSV infections and Helene Andrews-Polymenis for sharing the SL1344 *Salmonella* Typhimurium strain. We would like to acknowledge Andrew Hillhouse and the Institute for Genome Sciences and Society at TAMU for performing our RNA-sequencing experiments.

## MATERIALS AND METHODS

### Cell lines and bacterial strains

RAW 264.7 macrophages (ATCC) (originally isolated from male BALB/c mice) were cultured at 37°C with a humidified atmosphere of 5% CO_2_ in DMEM (Thermo Fisher) with 10% FBS (Sigma Aldrich) 0.5% HEPES (Thermo Fisher). For knockdown cell lines, RAW 264.7 macrophages were transduced with a pSICO-shRNA construct designed to target an exon or 3’UTR of an SR or hnRNP gene of interest. Knockdown macrophages were drug selected (hygromycin; Invitrogen) alongside a scramble (SCR) non-specific control. Each SR knockdown cell line was derived at the same time, as were the hnRNP cell lines. Knockdown efficiency of each factor was validated by RT-qPCR using exonic primer sets and the most efficient knockdown cell line (from 6 different knockdown constructs) was used for RNA-seq and the two most efficient knockdown cell lines were used for validation in Figs. 2 and 3.

### *S*. Typhimurium Infection

Infections with *Salmonella enterica* serovar Typhimurium were conducted by plating RAW 264.7 macrophages on tissue-cultured treated 12-well dishes at 7.5 ×10^5^ and incubated overnight. Overnight cultures of *S*. Typhimurium were diluted 1:20 in LB broth containing 0.3M NaCl and grown until they reached an OD600 of 0.9. Unless specified, cell lines at a confluency of 80% were infected with the *S*. Typhimurium strains at an MOI of 10 for 30 minutes in Hank’s buffered salt solution (HBSS). Infected monolayers were spun for 10 minutes at 1,000rpm, washed twice in HBSS containing 100μg/ml of gentamycin, and refreshed with media plus gentamicin (10 μg/ml). After removal of supernatant, cells were lysed in Trizol (Thermo Fisher) for RNA collection and analyzed using RT-qPCR.

### RNA-Seq

The RNA-Seq experiment was made up of 60 samples: biological triplicate of SCR uninfected, SCR *Salmonella*-infected, each knockdown uninfected, and each *Salmonella*-infected knockdown. RNA-Seq and library prep was performed by Texas A&M AgriLife Genomics and Bioinformatics Service. Samples were sequenced on Illumina 4000 using 2 × 150-bp paired-end reads. Raw reads were filtered and trimmed and Fastq data was mapped to the Mus musculus Reference genome (RefSeq) using CLC Genomics Workbench 8.0.1. Differential expression analyses were performed using CLC Genomics Workbench. Relative transcript expression was calculated by counting Reads Per Kilobase of exon model per Million mapped reads (RPKM). statistical significance was determined by the EDGE test via CLC Genomics Workbench. The differentially expressed genes were selected as those with p value threshold < 0.05 to include in the heatmaps represented.

### Gene Ontology (GO) Canonical Pathway and Hierarchical Analysis

To determine the most affected pathways in control versus knockdown RAW 264.7 macrophages, canonical pathway analysis was conducted using Ingenuity Pathway Analysis software from QIAGEN Bioinformatics. Genes that were differentially expressed wth a p value < 0.05 from our RNA-SEQ analysis were used as input from uninfected and Salmonella Typhimurium infected cells. Hierarchical clustering was done in Cluster3 (3.0) with complete linkage, absolute correlation (centered) parameters and visualized using Java TreeView.

### Scatter Plots and Correlation Analysis

For (p<0.05) differentially expressed genes, fold change was plotted to compare to coding sequence length which is identified by CLC Genomics Workbench to be equal to the total length of all exons (not all transcripts). Exon number and intron number were identified by CLC Genomics Workbench to be the number of exons/introns based on the mRNA annotations of the reference genome. Total gene length was calculated using “chromosome region start” and “chromosome region end” which are determined by CLC Genomics Workbench and the reference sequence to be the start position and end position of the annotated gene. Pearson Correlation was calculated using the values described above.

### RNA isolation and RT-qPCR analysis

For transcript analysis, cells were harvested in Trizol and RNA was isolated using Direct-zol RNA Miniprep kits (Zymo Research) with 1 hr DNase treatment. cDNA was synthesized with iScript cDNA Synthesis Kit (Bio-Rad). CDNA was diluted to 1:20 for each sample. A pool of cDNA from each treated or infected sample was used to make a 1:10 standard curve with each standard sample diluted 1:5 to produce a linear curve. RT-qPCR was performed using Power-Up SYBR Green Master Mix (Thermo Fisher) using a Quant Studio Flex 6 (Applied Biosystems). Samples were run in triplicate wells in a 96-well plate. Averages of the raw values were normalized to average values for the same sample with the control gene, *Actb*. To analyze fold induction, the average of the treated sample was divided by the untreated control sample, which was set at 1.

### Alternative Splicing Analysis

Alternative splicing events were analyzed using MAJIQ and VOILA with the default parameters [47]. Briefly, uniquely mapped, junction-spanning reads were used by MAJIQ to construct splice graphs for transcripts by using the RefSeq annotation supplemented with de-novo detected junctions. Here, de-novo refers to junctions that were not in the RefSeq transcriptome database but had sufficient evidence in the RNA-Seq data. The resulting gene splice graphs were analyzed for all identified local splice variations (LSVs). For every junction in each LSV, MAJIQ then quantified expected percent spliced in (PSI) value in control and knockdown samples and expected change in PSI (dPSI) between control and knockdown samples. Results from VOILA were then filtered for high confidence changing LSVs (whereby one or more junctions had at least a 95% probability of expected dPSI of at least an absolute value of 10 PSI units between control and knockdown) and candidate changing LSVs (95% probability, 10% dPSI). For the high confidence results (dPSI ≥ 10%), the events were further categorized as single exon cassette, multiexon cassette, alternative 5′ and/or 3′ splice site, or intron-retention.

### RBP Finder

For each gene, the canonical (longest) isoform of the gene (5’ and 3’ UTRs, plus CDS) as annotated by Ensembl (Mouse (GRCm38.p6)) was queried for SR/hnRNP motifs as defined by RBP-map. Stringency level was set on “High” and the Conservation Filter was applied. In cases where multiple motifs were listed, only a single “consensus” motif was chosen [55].

### VSV infection

7×10^5^ RAW cells were seeded in 12-well plates 16h before infection. Cells were infected with VSV-GFP virus [74] at multiplicity of infection (MOI) of 1 in serum-free DMEM (HyClone SH30022.01). After 1h of incubation with media containing virus, supernatant was removed, and fresh DMEM plus 10% FBS was added to each well. At indicated times post infection, cells were harvested with Trizol and prepared for RNA isolation.

### Quantitation and Statistical Analysis

Statistical analysis of data was performed using GraphPad Prism software. Two-tailed unpaired Student’s t tests were used for statistical analyses, and unless otherwise noted, all results are representative of at least three biological experiments (mean ± SEM (n = 3 per group)).

---

**Figure S1:**
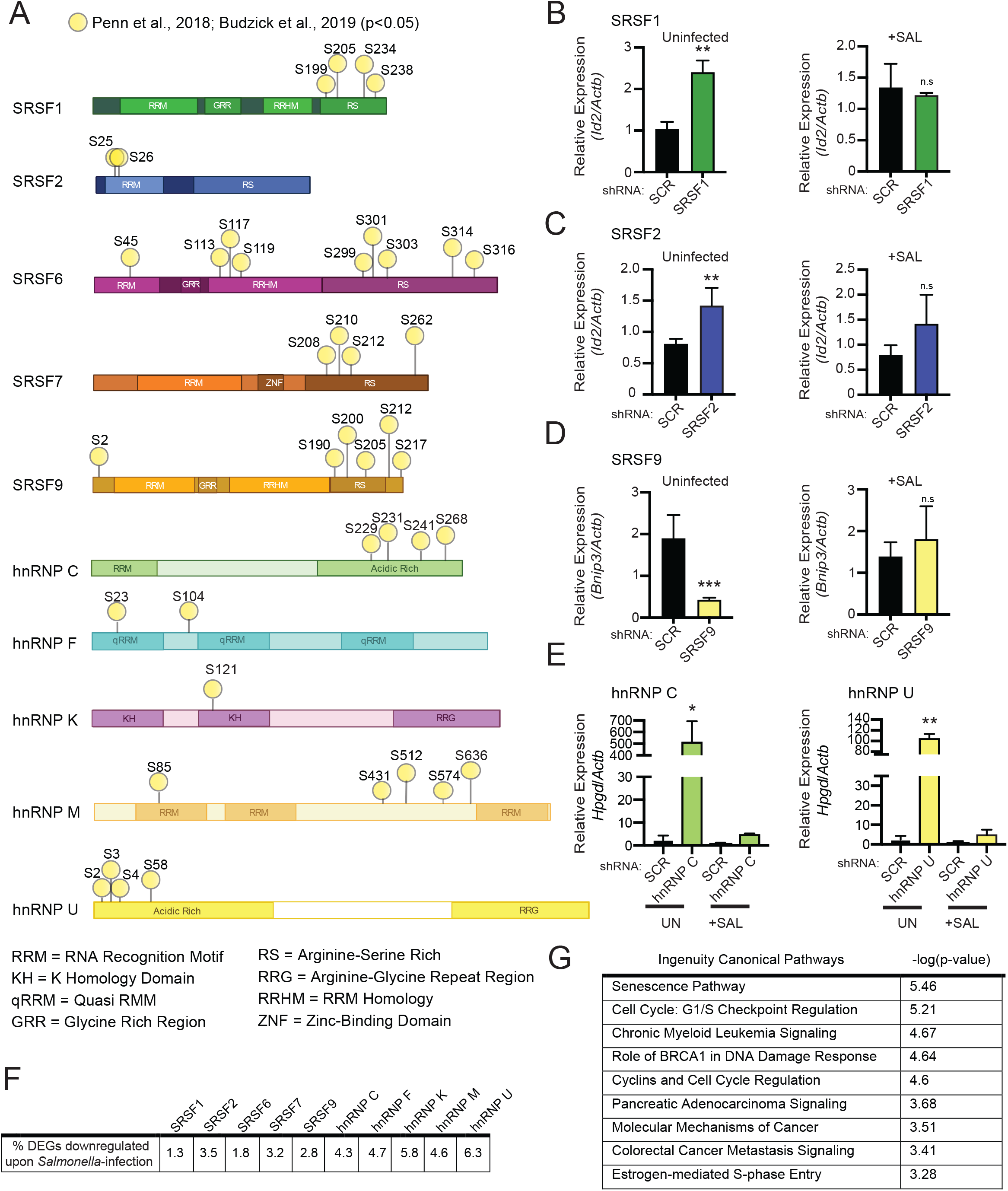
(A) Schematic representation of sites of differential phosphorylation reported by Penn et al., 2019 and Budzik et al., 2020 for SRSF and hnRNPs queried. Abbreviations for different protein domains are define below gene diagrams. (B) Relative gene expression of *Bnip3* over *Actb* in uninfected and *Salmonella*-infected SRSF1 knockdown macrophage cell lines. (C) As in (B) but *Id2* expression in SRSF2 knockdown macrophages. (D) As in (B) but *Bnip3* in SRSF9 knockdown macrophages. (E) As in (B) but *Hpgd* in hnRNP C and hnRNP U knockdown macrophages. (F) Percentage of differentially expressed genes (DEGs) in each SR/hnRNP knockdown cell line downregulated >2.0-fold in response to *Salmonella* infection in control macrophages. (G) Ingenuity pathway analysis (-log (p-value)) of genes downregulated in *Salmonella* vs. uninfected SCR macrophages (fold-change of -2 or more). For all RT-qPCRs, values are the mean of 3 biological replicates and error bars indicate standard deviation. * = p<0.05; ** = p<0.01; *** = p<0.005.

**Figure S2:**
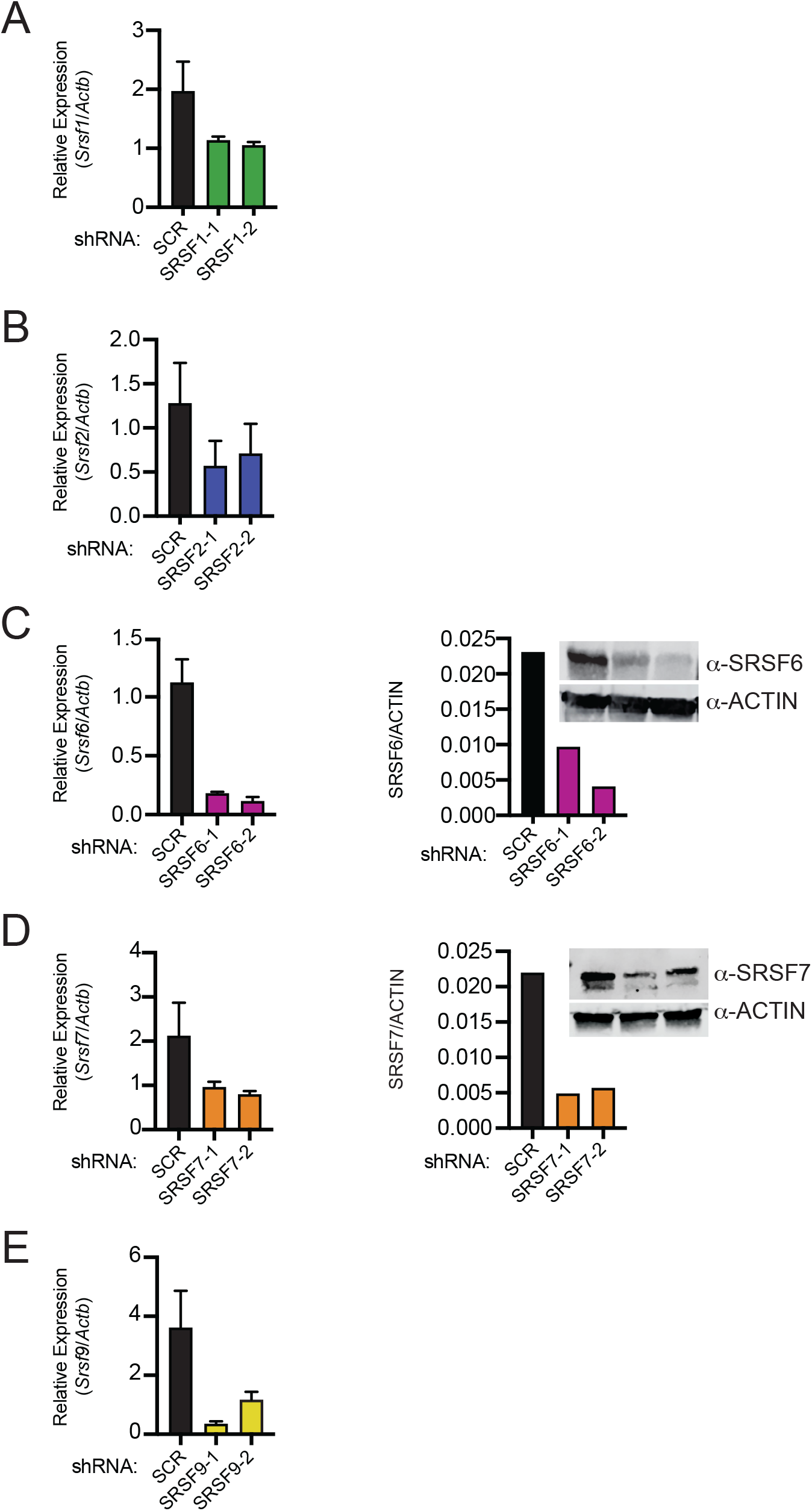
(A-E) Knockdown efficiency of SRSF RAW 264.7 macrophages by RT-qPCR (SRSF1, 2, 6 7, 9) and immunoblot (SRSF6, 7). For all RT-qPCRs, values are the mean of 3 biological replicates, error bars represent standard deviation, and immunoblots are representative of 2 or more independent experiments.

**Figure S3:**
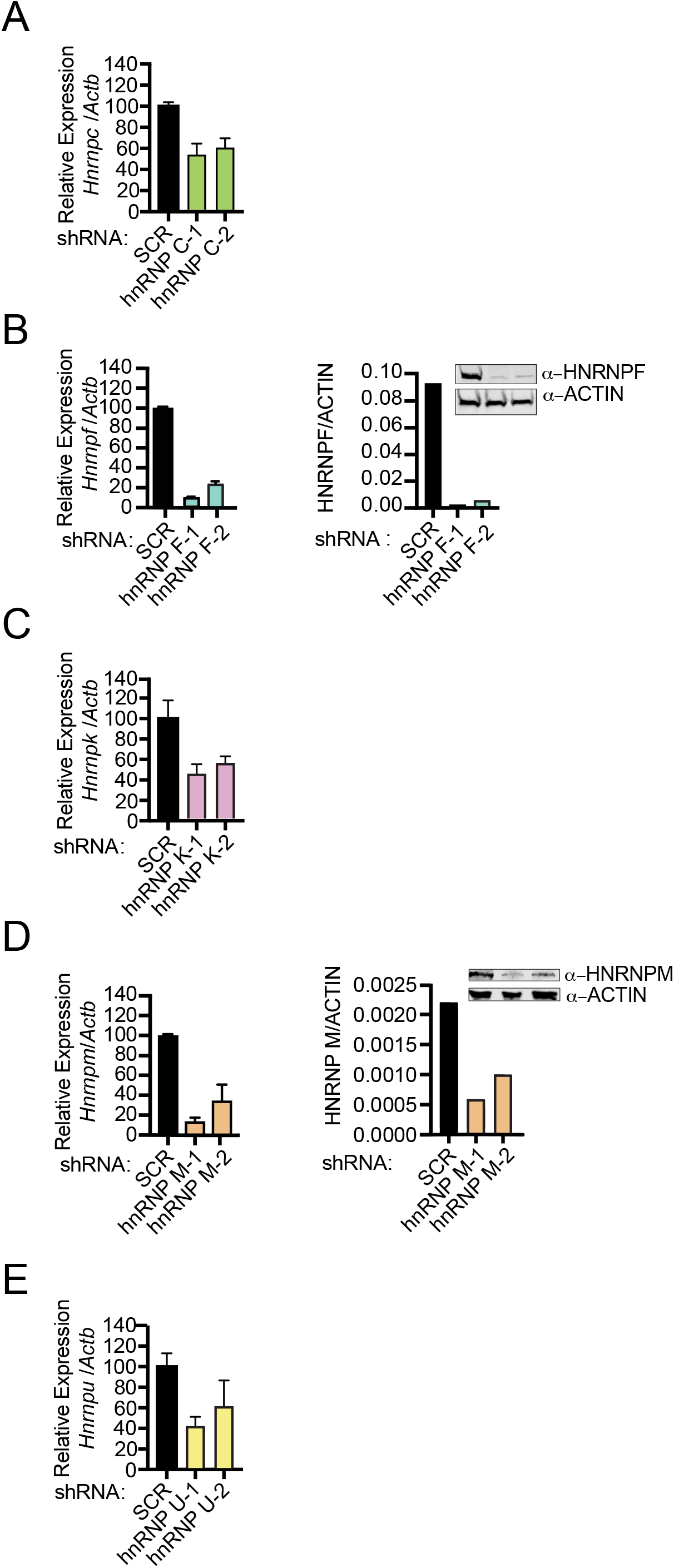
(A-E) Knockdown efficiency of hnRNP RAW 264.7 macrophages by RT-qPCR (hnRNP C, F, K, M, U) and immunoblot (hnRNP F, hnRNP M). For all RT-qPCRs, values are the mean of 3 biological replicates, error bars represent standard deviation, and immunoblots are representative of 2 or more independent experiments.

**Figure S4:**
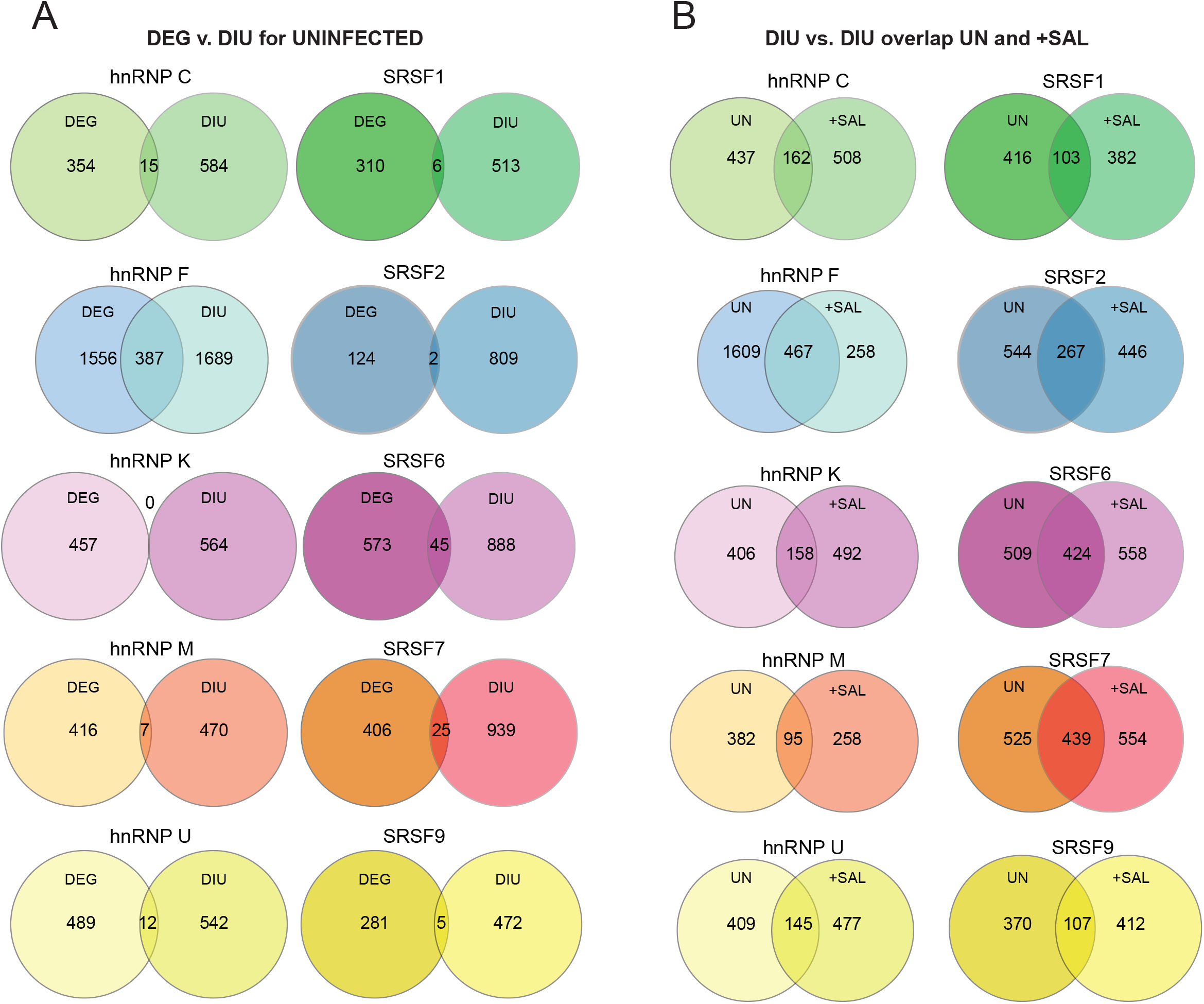
(A) Overlap of DEGs with genes that have significant MAJIQ LSVs (differential isoform usage (DIU) for hnRNP (left) and SRSF (right) knockdown RAW 264.7 macrophages vs. SCR controls. DEG p<0.05; LSV PSI (PSI ≥10%). (B) Overlap of genes with LSVs in uninfected vs. *Salmonella*-infected hnRNP (left) and SRSF (right) knockdown RAW 264.7 macrophages vs. SCR controls. LSV PSI (PSI ≥10%).

**Figure S5:**
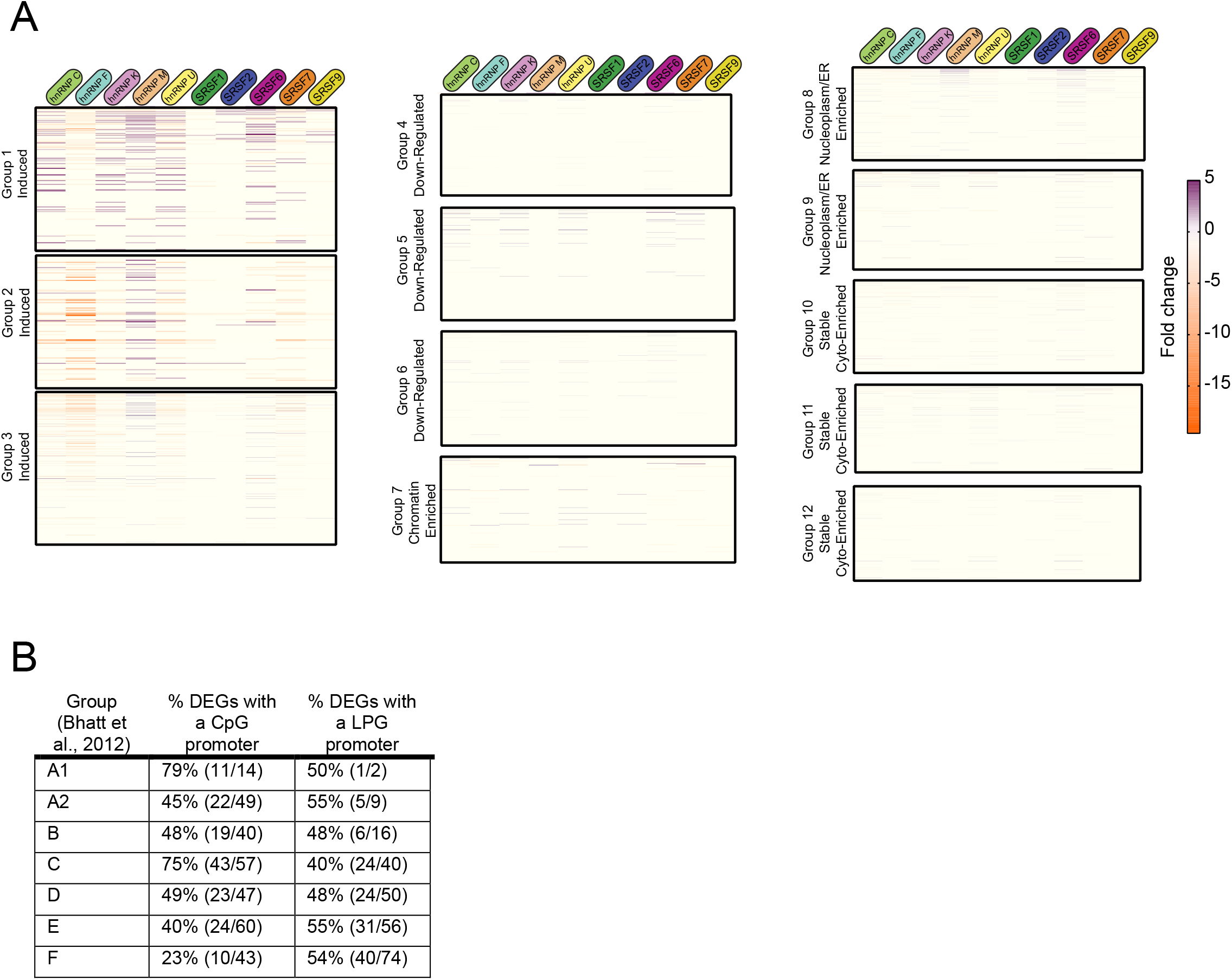
(A) Fold change of SR/hnRNP DEGs in *Salmonella*-infected RAW 264.7 macrophages compared to SCR controls with all genes (RefSeq genes exceeding 400-bp in length with an RPKM of at least 1 in at least one sample) categorized on basis of expression kinetics and subcellular accumulation in chromatin, nucleoplasm, and cytoplasm fractions as defined by Bhatt et al., 2012. (B) Calculation of genes with a high (CpG) and low CpG (LPG) promoter islands as defined by Bhatt et al., 2012 that were differentially expressed in one or more SR/hnRNP knockdown macrophage cell line compared to SCR control.

